# An investigation into the spatiotemporal patterns of the Nymphalid butterfly Vagrans egista sinha (Kollar, [1844])

**DOI:** 10.1101/2022.01.02.474748

**Authors:** Paul Pop, Kuldeep Singh Barwal, Randeep Singh, Puneet Pandey, Harminder Pal Singh, Sanjay Sondhi

## Abstract

Vagrans egista sinha (Kollar, [1844]), the Himalayan Vagrant is a subspecies of Nymphalid (Brush-footed) butterflies spread across Asia, whose western limit is in the north-west India. Observations of this subspecies have considerably increased over the past half-a-decade, with a spike in new sightings to the west of their previously known range. This has been considered as a range extension. The current study reports new records of this species from Bilaspur District, Himachal Pradesh, India (which are the first records for the district), through systematic and opportunistic sampling. This raises the question of whether the purported range extension towards the west could instead be a range shift or vagrancy, and whether there is any shift in elevational ranges in the populations across their known range. Questions pertaining to spatial differences in elevational ranges and seasonal variation, across their range, also piqued our curiosity. Using data from academic sources (such as published literature and museum collections), supplemented by data from public participation in scientific research and personal observations, these research questions are addressed. The accuracy of results when using citizen science data is also explored using the same dataset, focused on the impact of method of extraction of coordinates, and elevation derived from it under different scenarios. It was discovered that there has not been a range shift (either longitudinal or latitudinal) and observations don’t suggest vagrancy but a case of range extension. Other results indicated that there was no climb of population to higher elevations, no spatial differences in elevational ranges in the populations, or seasonal variation in activities across their range. It was also discovered that the method of data collection by, and extraction from, citizen science databases, can influence the accuracy of the results. Some problems involved in collecting data are discussed, and remedial solutions are suggested.

## Introduction

The genus Vagrans is considered as monotypic or bitypic by various authors (Kirti, Mehra & Sidhu, 2016). Vagrans egista sinha (Kollar, [1844]), the Himalayan Vagrant is a subspecies of Vagrans egista (Cramer, [1780]) - the Vagrant butterfly, a species whose distribution is far and wide in the Indo-Australian region, with the eastern limit being French Polynesia in the Pacific Ocean; western and northern limits at India; and the southern limit being Australia (GBIF, 2021). They are a largely orange-and-black coloured butterfly belonging to the Nymphalidae (Brush-footed) family, which is the largest family of butterflies (see Fig. 1 for the photograph of an individual). Different authors have adopted different genus, species, and subspecies names as well as author citations to refer to the Vagrant butterfly found in India — Vagrans sinha sinha Kollar, 1848 (Larsen, 2004), Issoria sinha sinha (Kollar) (Singh et al, 2011) and Vagrans sinha pallida Evans (Smetacek, 2012). We will be using the name Vagrans egista sinha (Kollar, [1844]), which is the most widely adopted name in circulation (Gokhale, 2021).

**Figure 1:**
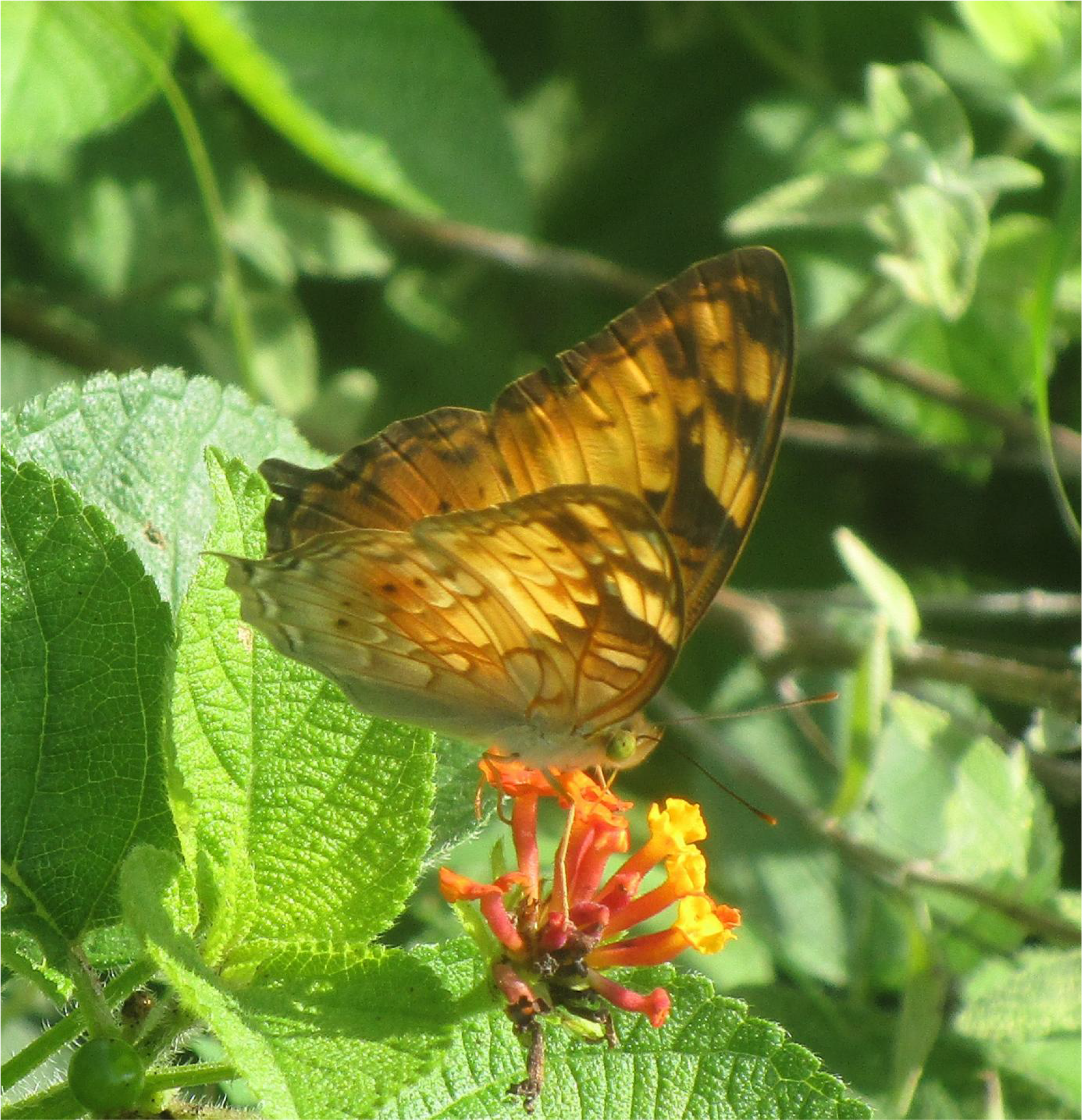
Species under study — Vagrans egista sinha (Kollar, [1844]), the Himalayan Vagrant

Their local occurrence varies from very common (Sondhi & Kunte, 2016) to rare (Bhowmik & Chowdhury, 2021), depending on the location. In India, this species is reported in tropical deciduous forests within 400-1400 m above sea level and Shorea robusta forests below 1000 m asl, and habitat-wise, they are found within riverine habitat, village homes near forest edges, and gardens (Smetacek, 2012; Sondhi & Kunte, 2018; Bhowmik & Chowdhury, 2021). Similarly, in Thailand and Vietnam, they are reported in very moist, lightly to partially shaded areas or gaps in the primary forest ( Spitzer et al, 1997, Küppers & Janikorn, 2009). The immature stages of this subspecies is recorded to feed on plants of the following genera: Dillenia (Dilleniaceae), Maytenus (Celastraceae), Flacourtia, Homalium, and Xylosma (all Salicaceae members) (compiled in Gokhale & Yathumon, 2021). The same study reports the confirmed use of Xylosma longifolia Clos by this species as a larval host plant in Dehradun (Western Himalaya). The adults are observed to feed on flower nectar (Polygonum hydropiper L. - Water Pepper Plant and Lantana camara L. - Wild Sage, being two such plants) and animal excretions (bird droppings, cow urine and dung), and also involve in mud-puddling (Naro & Sondhi, 2014; Sondhi & Kunte, 2018; Sharma & Sharma, 2018; Kumar, Kushwaha & Namdev, 2020). They have been found to opportunistically feed on flower nectar in the forest canopy during some migrations in the Tam Dao mountains in Vietnam (Spitzer et al, 1993). Migrations are not recorded in India. Parsons & Cantlie (1948) report that they are known to sit on hot tarmac and gravel by the roadside in winter, most likely an adaptation to warm up during the cold season. They also hint at possible variation in size in Assam compared to adjacent areas.

This subspecies is distributed in 10 countries (earliest known explicitly mentioned record given in brackets) (Fig. 2): India (Bingham, 1905), containing the most northern and western distribution of the species as well as the subspecies; Nepal (Varshney, 1994); Bhutan (Singh, 2012); Bangladesh (Larsen, 2004); Myanmar (Bingham, 1905); Thailand (Varshney, 1994), containing the southernmost distribution of the subspecies; Cambodia (Savela, nd); Laos (No record, but most likely); Vietnam (Spitzer et al, 1993); and China (D’Abrera, 1940), including the special autonomous region Hong Kong (Bascombe, Johnston & Bascombe, 1999) — containing the easternmost distribution of the subspecies. In India, they are found within 14 states and one union territory across the Himalayan range, its foothills, Eastern Ghats and near the Eastern coast — Jammu & Kashmir (Sharma & Sharma, 2018), which is the northernmost and westernmost distribution in their range; Himachal Pradesh (Kirti, Mehra & Sidhu, 2016); Chandigarh (Soman, 2015); Uttarakhand (Bingham, 1905); Uttar Pradesh (Kumar, Kushwaha & Namdev, 2020); Sikkim (Wynter-Blyth, 1957); West Bengal (Bingham, 1905); Arunachal Pradesh (Kunte, 2007), their easternmost distribution in India; Assam (Bingham, 1905); Nagaland (Tytler, 1911); Mizoram (Bhakare, 2009); Tripura (Zothansangi, 2015); Odisha (Wynter-Blyth, 1957); Chattisgarh (Naidu, 2018); and Andhra Pradesh (Saragada, 2020), which is their southernmost distribution within India.

**Figure 2:**
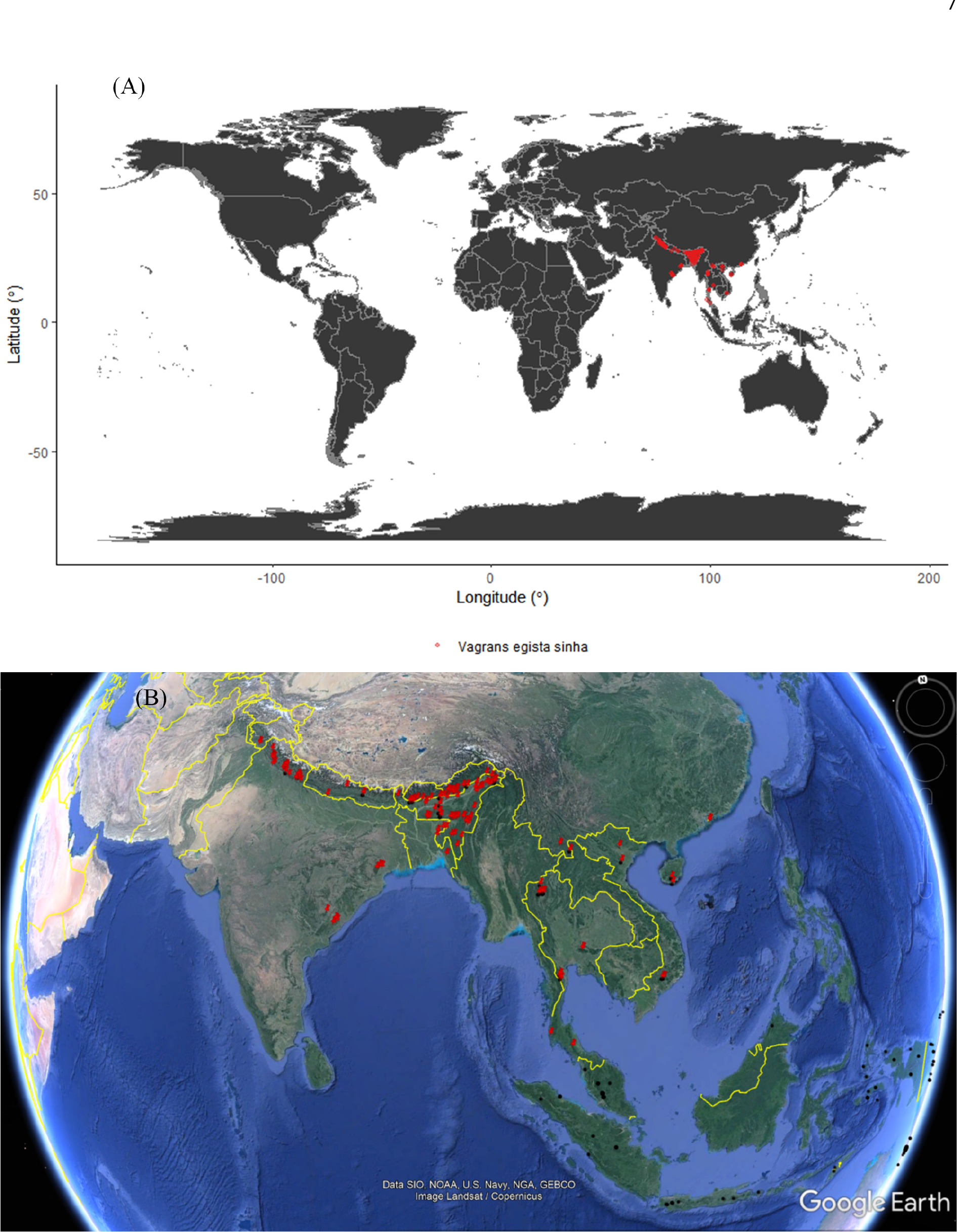
(A) Global distribution of the subspecies on a WGS 1984 projection of Earth. (B) Individual records of the subspecies (red pins) and species (black circles) on a globe.

Records of this subspecies have increased considerably over the past half-a-decade, with many new sightings to the west of their previously known range in North-West India. Upon cursory glance, there is also a lack of recent records in much of South-East. This apparent range extension towards the west could therefore be a range shift or a case of vagrancy. This may indicate disturbance in parts of their range or simply processes such as increasing population sizes, overshoot of dispersal range or getting blown off by wind. We adopt Jain et al’s (2018) definition of vagrancy as sporadic sightings of a species with no breeding records. While Jain et al (2018) definition allowed for a maximum of three individuals in the span of 17 years in a country, we modify it and adopt the following criteria, failing any of which a species cannot be considered as a vagrant: i) maximum of 3 sightings per year in a geo- political unit, which is a district in our case (which is far less than for a migratory species of butterfly) ii) no breeding records iii) no repeated sighting in the same unit (district) in atleast two consecutive years. Range shift is defined as the changes in geographic dimensions of a species range (which are latitude, longitude, and elevation) over the fourth dimension i.e. time (Lenoir & Svenning, 2013). For analytical purposes, this indicates the presence of a leading edge with expansion into previously unoccupied area, as well as a trailing edge where the species is going locally extinct (Lenoir & Svenning, 2013). Range extension is equal to range shift minus trailing edge. This can confirmed by default, if there is no evidence for range shift or vagrancy.

“Citizen science” refers to public participation in scientific research (‘c-science’ henceforth). C-science is extensively used in lepidopterology, especially for butterflies, with the number of species studied and sites monitored outweighing all vertebrate groups in Europe (Schmeller et al, 2009). It is used for several purposes, such as collecting photographs of rare butterflies (Mesaglio et al, 2021), monitoring populations at a national level (Sanderson, Braby & Bond, 2021), and even studying the range expansion of a Nymphalid butterfly species distributed through the Indomalayan and Australasian realms (similar to the current study) (Chowdhury et al, 2021). Reliability of c-science data, especially when it comes to spatiotemporal patterns of butterflies depends on a wide-variety of components of a c-science model, including accuracy of identification of species by participants, particularly for cryptic species (Vantieghem et al, 2016); irregular nature of survey efforts over time; incompleteness and selectivity in observation; geographical bias (van Strien, van Swaay, & Termaat, 2013).

Studies of butterflies have been done to compare the reliability of c-science data (opportunistic data) in comparison to standardized monitoring data. In assessment of distribution trends using occupancy models (van Strien, van Swaay, & Termaat, 2013), and species population trends (Dennis et al, 2017), it was shown that the former performs almost as well as the latter. Differences have been found in the comparison of quantity and quality of outputs of two regions (Matteson, Taron & Minor, 2012) when the monitoring protocols are different: c-science resulted in more species due to unrestricted sampling but standardized monitoring were more likely to detect trends due to consistent sampling. While much work has been done on the reliability of c-science data, as far as we know, there has been no study to look at the influence of reliability of coordinate data derived through various means, and multiple scenarios of secondary data derived from these coordinates (such as elevation) to be used as variables/covariates.

We addressed five questions, namely: 1) are increasing records of the subspecies westwards of their known range indicative of a range extension, range shift, or vagrancy?; 2) are there seasonal changes in distribution across its range?; 3) are populations shifting to higher elevations across their known range, possibly indicative of climate change?; 4) is there a gradient of elevation across distributions of populations; and 5) how reliable is c-science data for addressing such ecological questions? Questions 1 and 3 are prespecified (hypothesis-based) whereas the rest are exploratory in nature (some subsections within the prespecified questions were also exploratory).

## Materials and Methods

### Data characteristics

As part of the project “High Resolution Spatial Mapping of Bird Phenology as an Indicator of Ecosystem Health in Relation to Climate Change in Himalaya”, surveys (which includes invertebrate monitoring) were undertaken in Bilaspur District of Himachal Pradesh from March 2020 to April 2021, through which some of the data were collected. These surveys included both reconnaissance surveys (done to design a stratified random sampling protocol) and stratified random sampling points (designed for systematic surveys of birds, invertebrates and plants). Identification was carried out using Sondhi & Kunte (2018), an easy task as there are no species that are confusingly similar in the region. Bulk of data were collected for the current study from a c-science website (Butterflies of India); a data aggregator (GBIF, which contained data from c-science portals such as iNaturalist and India Biodiversity Portal, as well as records of specimen collections in museums); peer-reviewed and grey literature; and communication with authors of the literature. A total of 216 usable data points were obtained from this (Fig. S1) for a total of seven countries (Table S1; a record from one country — Cambodia was missed at the time of analysis, and hence excluded), the contribution from literature and communication with authors constituting the bulk at ∼55.1%, followed by c-science at ∼36.1% . Of the 24 authors contacted (including multiple authors of the same paper) through email or websites such as Research Gate and Academia Edu regarding 20 papers for more information regarding their records, only nine people responded (37.5% response rate) regarding 10 papers (50%) covering various countries such as India (Singh, Gupta & Varatharajan, 2011), Bangladesh (Khandokar et al, 2013), China (Kendrick (Ed), 2005), and Thailand (Küppers & Janikorn, 2020). All except one was through email (one being Research Gate messaging). Coordinates for location names were deduced using the given landmarks, elevation information, given coordinates of adjacent locations, comparison between historical names and current names etc. For the current study, no records were available to a desired resolution needed for use in analysis from Myanmar, Laos and Cambodia.

### Data analysis

Data sorting, analysis and visualisation were carried out using WPS Excel, R (v 3.6.1), R Studio (v 4.1.0), and QGIS (v 3.8.2) for studying the autecology of the V. e. sinha. Selection: Some records were omitted due to locations covering a large area and ambiguity in their name (due to recent changes in the name). In one case where there were two records within half an hour interval on the same site, only one record was retained. When there were entries duplicated in multiple sources, only the one with the most reliable information (such as closeness of name of location and coordinates provided) was retained. For sight-based records with no coordinate uncertainty information, different coordinate uncertainty values were assigned for analysis, ranging from 0 km to 10 km. For museum specimens with no coordinate uncertainty information, at least 5 km was assumed unless more details were available. The assignment of coordinate uncertainty was carried out under manual supervision, based on all the available information for each record (such as coordinates within an area such as a National Park having a coordinate uncertainty equal to the radius from the point that which could be realistically covered by walking or in a vehicle depending on the availability of trails or roads, whose limit is set by the boundaries of the Park).

To retain a greater number of records, an novel approach has been taken wherein median dates were calculated, as well as adjusted-median dates. Median calculation was done as follows: When date range was only mm-yyyy, dd was assumed to be 01 (first day of that month) for begin dates and 28 to 31 (last day of that month) for end dates. Similarly, for yyyy, dd-mm were assumed to be 01-01 (first day of that year) for begin dates and 31-12 (last day of that year) for end dates. For dates with only one mm-yyyy, dd was assumed to be 14-16 (mid-month) and similarly for only one yyyy, dd-mm were assumed to be 01-07 (mid-year). Median date was also adjusted for the information about the months reported. For example, if the study was done from March 2012 to February 2013, the median date would be the 30^th^ of August 2012. If it was given that the species was recorded from September-October, then the date was adjusted to the closest month i.e. the 30^th^ September, 2012. This is the adjusted- median date. Dates were converted to date values to use as a continuous measurement. Since individual zones can have different patterns, three clusters were compared using histograms: the Western Himalayan cluster, including the foothills; the North-East Indian cluster, including Bhutan and Bangladesh; and the South-East Asian cluster, including China.

Data were tested for normality, using the Shapiro-Wilk test and q-q plots (values from Shapiro-Wilk test and q-q plots are available in Table S2 and Fig. S2-S5, respectively). Shapiro-Wilk test is inadvisable when sample sizes are extremely large, which as a rule of thumb is >250 (Ruxton, Wilkinson & Neuhäuser, 2015). As our data were a maximum of 216 datapoints, it was used. However q-q plots were used to make the final decision. When the p-value was <0.05, the null hypothesis that the data are normally distributed was rejected (alpha level = 0.05). If the p-value was > 0.05, then the null hypothesis was not rejected. For hypothesis testing using correlation, it was decided that Pearson’s Product-Moment Correlation (PPMC) would be used if the data were normal, and if non-normal, Spearman’s (rho) and Kendall’s rank correlation (tau) were to be used (Crichton, 2001). Spearman’s rho was ignored and Kendall’s tau was given importance when the used data had a considerable amount of ranking ties because tau is more accurate in that condition (Nie, Bent & Hull, 1970). The statistical interpretation of PPMC and rho were based on Dancy & Reidy (2007) (+/−1 = perfect correlation, +/−0.9 to +/−0.7 = strong, +/−0.6 to +/−0.4 = moderate, +/−0.3 to +/−0.1 = weak, and 0 = zero), which is a rather conservative scale compared to other approaches (see Akoglu, 2018). Tau was interpreted as follows: +/−1 = perfect correlation, +/− 0.99 to +/− 0.40 = strong, +/−0.39 to +/−0.35 = moderate, and +/−0.34 to +/−0.1 = weak, and 0 = zero. Final decision was made based on the context of questions addressed, as is generally advised (Schober, Boer & Schwarte, 2018). To avoid p-hacking (Head et al, 2015), the tests were carried out under different scenarios, including outliers removed, and p-values for all these tests were reported. To avoid overemphasis on p-values due to its rather unreliable nature (Nuzzo, 2014), visualisation of data was carried out when necessary and results were interpreted with biological plausibility and relevance in consideration. Histograms were created for all the variables considered.

For addressing the first question, check for negative correlations between longitude and dates or years (which indicates westward shift) were done, and observations were checked for fulfillment of the criteria for vagrancy. For understanding seasonal variation, correlations between months and coordinates were done. Third, the question was addressed by checking for a positive correlation between elevation and dates. The exploratory analysis to answer the fourth question involved checking for positive or negative correlations between elevation and coordinates.

The final question was tackled using the following steps. To find out the differences between coordinate deduced from name and that directly acquired from the source, 39 unique datapoints from the Butterflies of India website (Gokhale, 2021) were used to find the shortest distance between them and 39 corresponding acquired ones. The distance was found using the Vincenty Ellipsoid method which is accurate up to a few millimetres, using the R package geosphere. A subsample of these distances was manually verified for accuracy using Google Earth. For points with no reported coordinate uncertainty (a total of 63 points, a majority of them from c-science sources), four scenarios were considered. In the first scenario, the coordinates reported were assumed to be completely accurate. In the second, third and fourth scenario, buffers of 0.5, 5, and 10 km were drawn with the coordinates as the centre, and the mean elevation for each of such coordinates was derived from the ASTER GDEM raster layer (ASTER GDEM is a product of METI & NASA) using zonal statistics plugin in QGIS (Fig. S6). Note that due to the 2D nature of the buffer and 3D nature of the Earth, the fixed-width buffer actually had variable length from the centre, which is offset by the fact that this variation was present across the range. The biological relevance of this step is that it calculates an elevation where the subspecies probabilistically has a higher chance of being for the largest amount of time. Even in cases where the elevation of a record is known to 1 m accuracy, buffers like these give a more biologically realistic idea of the elevation due to the motile nature of butterflies. All such scenarios were compared by checking for correlation with other variables, by comparing the correlations between scenario 1 and scenarios 2-4 and, as well as doing a one-way ANOVA. For the latter two steps, 63 points alone as well as these points included in the 216 total points were analysed separately to understand the variations of the values of variable buffer points alone, as well as their influence on data when mixed with unchanging elevation values. Tests to satisfy the assumptions of ANOVA were also carried (independence was assumed, qq plots for normality, and Bartlett’s test for homogeneity of variance).

## Results

During field surveys, five sightings of V.e.sinha in their adult stage were made in the district (see Fig. 3 for sighting details). Two of these sightings were made during reconnaissance surveys and three of these sightings were made within stratified random sampling points. In Rani Kota, two individuals were sighted, whereas in Kiarian, three individuals were sighted, two of them likely mating in the air near and on a Mangifera indica (Mango) tree. At rest of the sighting localities, only one individual was found. In a majority of the sightings, they were feeding on the invasive plant L. camara (Fig. 1), a behaviour which has also been seen in Uttar Pradesh (Kumar, Kushwaha & Namdev, 2020).

**Figure 3:**
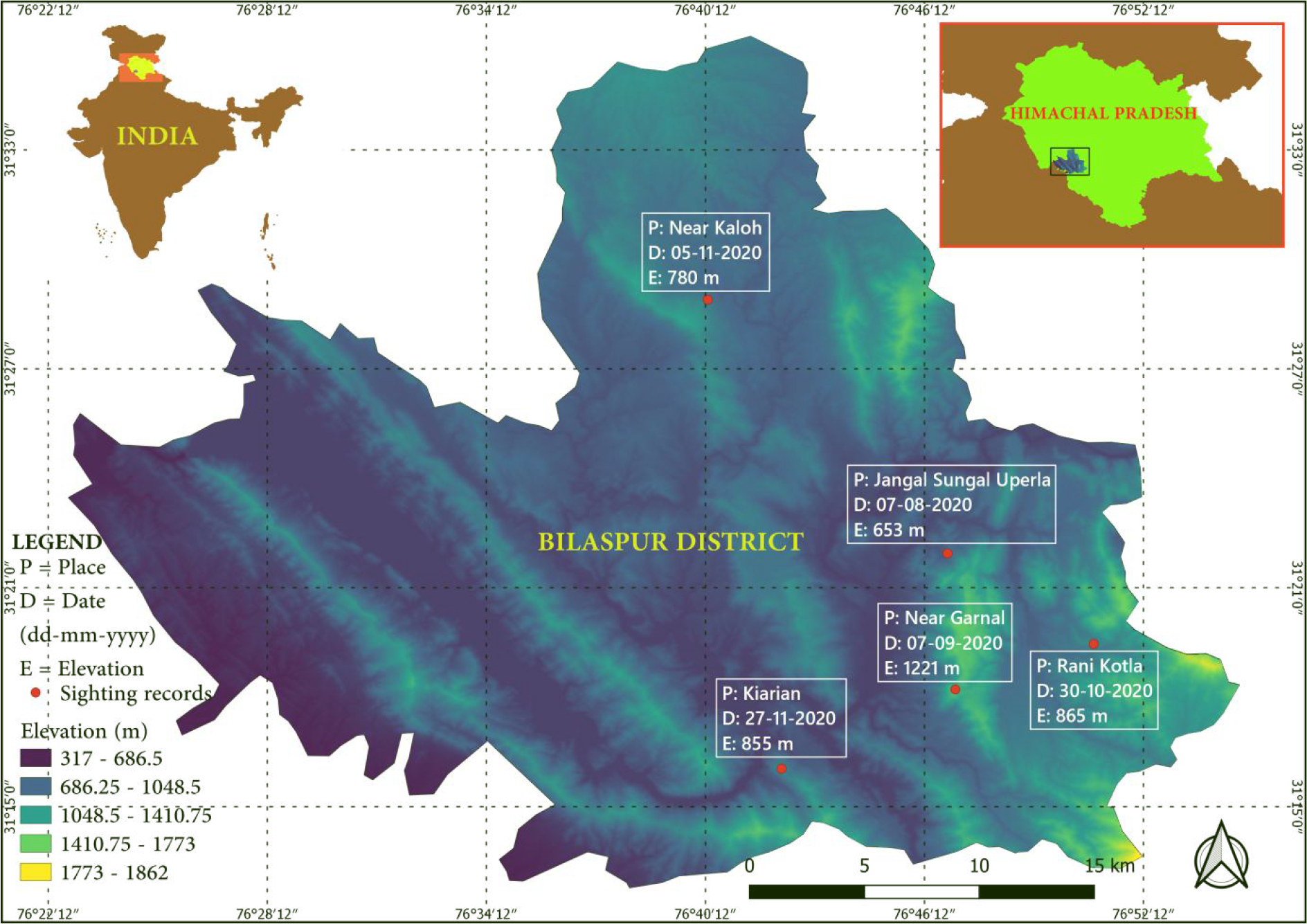
Sightings of the species during the current study from Bilaspur District, Himachal Pradesh.

Date, latitude and longitude were found to be non-normal data in all scenarios, except for elevation which was found to follow a normal distribution. Hence, non-parametric tests were used for most cases. Rho and tau were used for most of the scenarios, except for addressing the question on the reliability of c-science data, where PPMC was used. It can be observed from the histogram of years (Fig. 4E), that most of the records collected were after 2010, a lesser proportion between 2000 and 2010, and even lesser from before the start of the 21^st^ century, indicating a skew towards the last two decades. When histograms were generated cluster-wise (Fig. 5A), it has been found that most records from earlier years were from the SE Asian cluster, and over there, records in recent history are comparatively few. In the NE Indian and the W Himalayan clusters, records started growing from 2008, with the former having tall peaks during the last decade, whereas the latter showed a relatively consistent and moderate number of records. Month-wise distribution of records show a bimodal distribution nature with a small peak at spring/post-winter and a taller and wider peak during the final months of monsoon and most of the winter/post-monsoon months (Fig. 4C). Individual histograms for the three clusters (Fig. 5B) indicated bimodal distribution in all the clusters.

**Figure 4:**
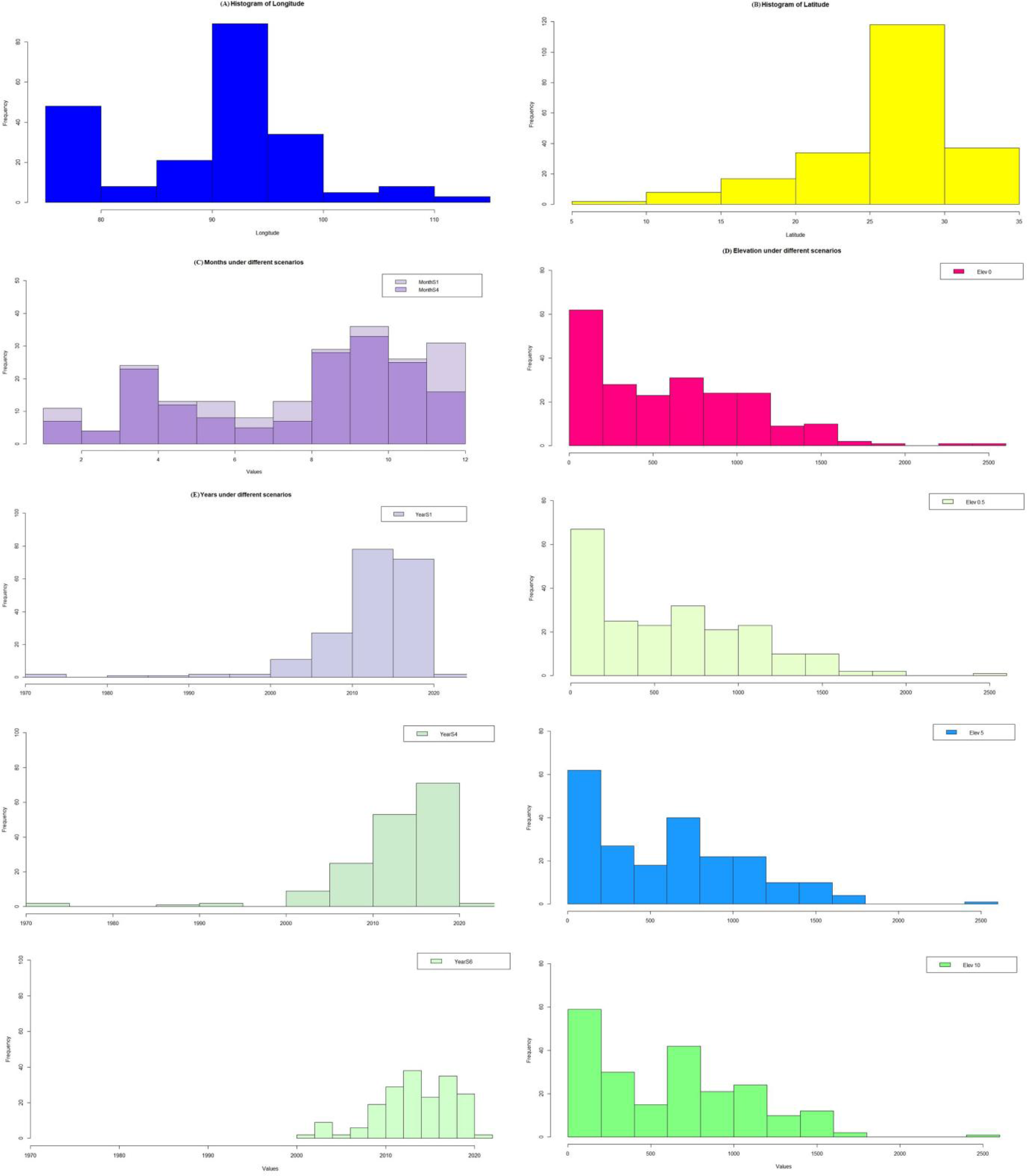
Histogram of variables across the species range. (A) Histogram of longitude (n = 216). (B) Histogram of latitude (n = 216). (C) Months under different scenarios — S4 being actual months (n = 168) and S1 being actual+median+adjusted median dates (n = 208). (D) Elevation under different scenarios (n = 216, four figures in the same column) — Elev 0, Elev 0.5, Elev 5, Elev 10 (with 0 km coordinate uncertainty kept as zero or changed to 0.5, 5, and 10 km respectively, for 63 points). (E) Years under different scenarios (three figures in the same column) — Year S1, Year S4, and Year S6 (YearS1 = actual years + median years + adjusted median years, n = 198; YearS4 = actual years, n = 165; and Year S6 = YearS1 - pre- 2000 records, n = 190).

**Figure 5:**
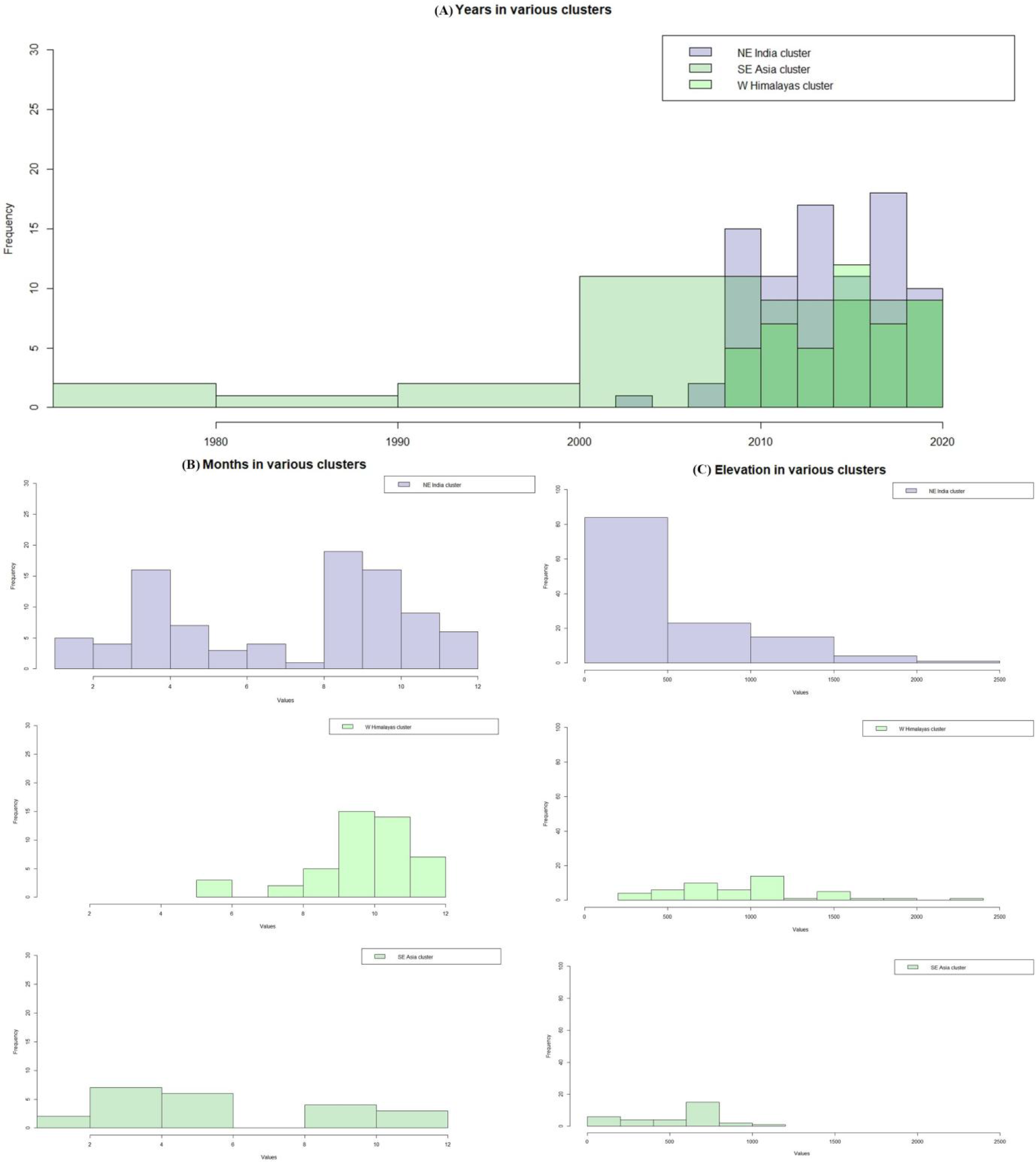
Histograms of variables in different clusters — the North-West Himalayan cluster, the North-East Indian cluster, and the South-East Asian cluster. (A) Years in various clusters, combined. Please note that in areas of overlap, colours are additive. In areas where the three clusters overlap, colour is bright dark green; Indian clusters together = medium green; and NE Indian+SE Asian = purplish-green. (B) Months in various clusters. (C) Elevation in various clusters.

However due to data deficiency and spread in the SE Asian cluster, it cannot be asserted with confidence that this is the pattern for this area.

The elevation histogram (Fig. 4D) shows two peaks, indicating most sightings at the elevation between 0 and 200 m asl and around half that many between 600-800 m asl. Individual histograms (Fig. 5C) show that this pattern is mostly contributed by records from the NE Indian cluster and the W Himalayan cluster which show the largest peak from 1000-1200 m asl (but could be an artefact of sampling bias). In the longitude histogram (Fig. 4A), there are two major uphills — between 75-80° E being Uttarakhand, Himachal Pradesh, and Jammu & Kashmir (most records from Uttarakhand), and between 90-95° E being North-East India, and a small adjacent peak from 95-100° E showing records from Thailand. Latitudinal distribution (Fig. 4B) shows a marked high frequency between 25-30° N, which is where North-East India and parts of Uttarakhand (Nainital District) are located.

Main findings are as follows:

1. No significant positive or negative linear relationships between dates (as absolute dates and years) and coordinates were found, indicating no range shift, either longitudinal or latitudinal. The results of this are summarized in Table 1 and 2. Note that in scenario 5 for correlations between coordinates and years, the p-value was > 0.05. This indicates that the null hypothesis that these two variables are uncorrelated is not rejected. However, this may be due to low sample sizes. This scenario is the least realistic as the actual known dates are removed. In scenario 6 and 7 which are realistic ones, where the sparse (outlier) data from pre-2000 and pre-2005 have been removed, the p-value was almost always >0.05, indicating non- correlation. Although vagrancy seems like a plausible explanation for the sightings of the subspecies in Himachal Pradesh and Jammu & Kashmir, it is unlikely due to repeated sightings in Solan District across years and the relatively high number of observations in Bilaspur District in a multi-season multi-year study (March 2020 to April 2021). Surveys were often single-season or single-year, and not multi-season multi-year. This sampling bias has resulted in knowledge gaps. The recent observations in their north-western-most part is, therefore, most likely a case of range extension.
2. No significant correlations between months and coordinates were found, indicating no linear pattern of seasonal variation (summarised in Table 3). However, Spearman’s correlation between Longitude and Months showed a moderate negative relationship. In this case, Tau was given priority due to a considerable amounts of tied ranks. Like earlier, probably as a result of a lower sample size, scenario 5 showed p-values of > 0.05.
3. No significant linear correlations between elevation and date were found, indicating that no climbing up to higher elevations occurred over the years (Table 4). P-values never dropped below 0.05 but were consistently high. Hence they are likely to be highly uncorrelated.
4. No significant linear correlations between latitude and elevation, or longitude and elevation, were found, indicating no linear pattern of elevation across populations (Table 5).
5. Using the Vincenty Ellipsoid method, the mean shortest (straight line) difference in distances between corresponding pairs of points was found to be 7855.95 m (Median = m, St dev (pop) = 9475.03 m, Range = 0.298 - 38809.5 m). The correlation between elevations under different scenarios have shown that there is a linear decline in correlation between coordinates with fixed or no buffers versus coordinates that had changing buffers, both individually and when mixed with coordinates of fixed buffer values (Table 6).

**Table 1:**
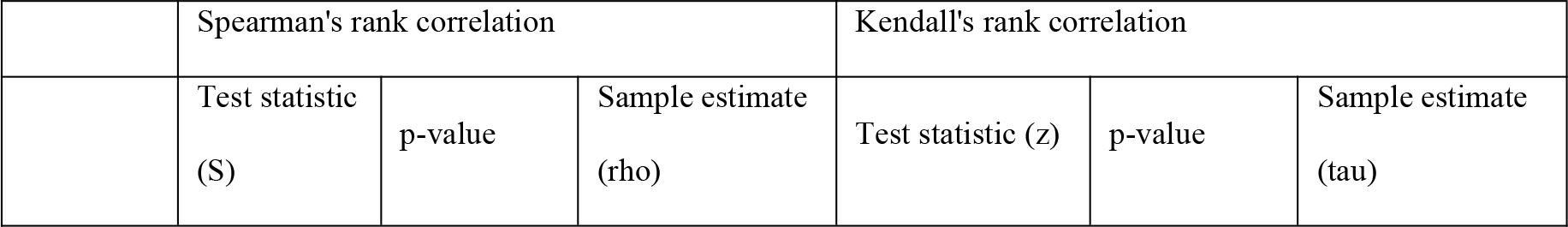
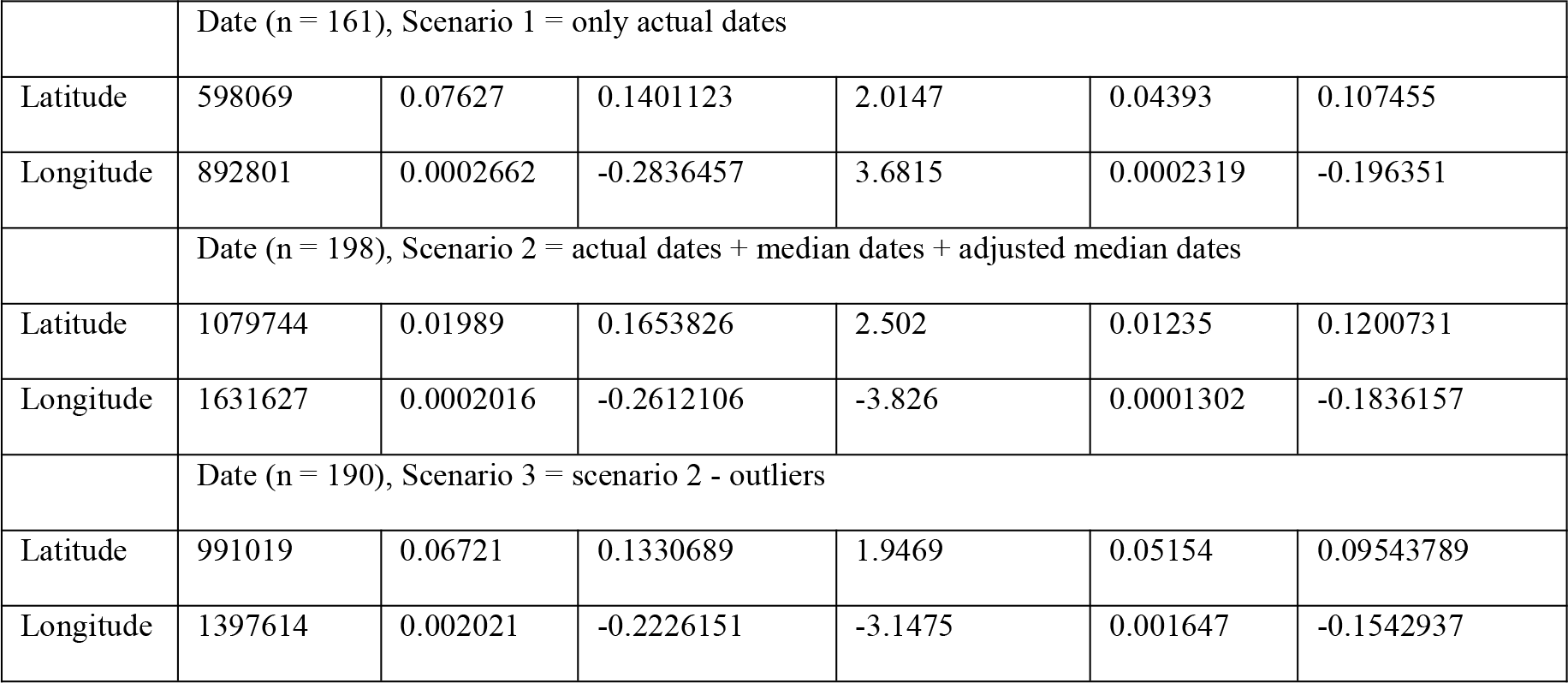
Correlations between dates and coordinates.

| Spearman's rank correlation |  |  | Kendall's rank correlation |  |  |
| --- | --- | --- | --- | --- | --- |
| Test statistic<br>(S) | p-value | Sample estimate<br>(rho) | Test statistic (z) | p-value | Sample estimate<br>(tau) |

|  |  |  |  |  |  |  |
| --- | --- | --- | --- | --- | --- | --- |
|  | Date (n = 161), Scenario 1 = only actual dates |  |  |  |  |  |
| Latitude | 598069 | 0.07627 | 0.1401123 | 2.0147 | 0.04393 | 0.107455 |
| Longitude | 892801 | 0.0002662 | -0.2836457 | 3.6815 | 0.0002319 | -0.196351 |
|  | Date (n = 198), Scenario 2 = actual dates + median dates + adjusted median dates |  |  |  |  |  |
| Latitude | 1079744 | 0.01989 | 0.1653826 | 2.502 | 0.01235 | 0.1200731 |
| Longitude | 1631627 | 0.0002016 | -0.2612106 | -3.826 | 0.0001302 | -0.1836157 |
|  | Date (n = 190), Scenario 3 = scenario 2 - outliers |  |  |  |  |  |
| Latitude | 991019 | 0.06721 | 0.1330689 | 1.9469 | 0.05154 | 0.09543789 |
| Longitude | 1397614 | 0.002021 | -0.2226151 | -3.1475 | 0.001647 | -0.1542937 |

**Table 2:** Correlations between years and coordinates.

|  |  |  |  |  |  |  |
| --- | --- | --- | --- | --- | --- | --- |
|  | Spearman's rank correlation |  |  | Kendall's rank correlation |  |  |
|  | Test statistic (S) | p-value | Sample estimate (rho) | Test statistic (z) | p-value | Sample estimate (tau) |
|  | Year (n = 198), Scenario 1 = actual years + median years + adjusted median years |  |  |  |  |  |
| Latitude | 1094096 | 0.02998 | 0.1542888 | 2.3691 | 0.01783 | 0.1170505 |
| Longitude | 1606976 | 0.0005879 | -0.2421557 | -3.5747 | 0.0003507 | -0.1766173 |
|  | Year (n = 195), Scenario 2 = actual years + median years |  |  |  |  |  |
| Latitude | 1058328 | 0.04521 | 0.1435955 | 2.1907 | 0.02848 | 0.1091555 |
| Longitude | 1516382 | 0.001412 | -0.2270651 | -3.3009 | 0.0009638 | -0.1644785 |
|  | Year (n = 168), Scenario 3 = actual years + adjusted median years |  |  |  |  |  |
| Latitude | 664048 | 0.03867 | 0.159693 | 2.3041 | 0.02122 | 0.1240884 |
| Longitude | 1016653 | 0.0001665 | -0.2865052 | -3.8052 | 0.0001417 | -0.204935 |
|  | Year (n = 165), Scenario 4 = actual years |  |  |  |  |  |
| Latitude | 639561 | 0.06181 | 0.1457253 | 2.0872 | 0.03687 | 0.1135471 |
| Longitude | 950502 | 0.000462 | -0.2696045 | -3.5193 | 0.0004327 | -0.1914519 |
|  | Year (n = 33), Scenario 5 = median years + adjusted median years |  |  |  |  |  |
| Latitude | 7597.9 | 0.129 | -0.2697014 | -1.7569 | 0.07893 | -0.2331204 |
| Longitude | 5614.3 | 0.7327 | 0.06178231 | 0.11495 | 0.9085 | 0.01526533 |
|  | Year (n = 190), Scenario 6 = Scenario 1 - pre-2000 records |  |  |  |  |  |
| Latitude | 1005034 | 0.09685 | 0.1208094 | 1.804 | 0.07123 | 0.09126776 |
| Longitude | 1373053 | 0.005393 | -0.2011293 | -2.8784 | 0.003997 | -0.1456269 |
|  | Year (n = 177), Scenario 7 = Scenario 1 - pre-2005 records |  |  |  |  |  |
| Latitude | 874965 | 0.4815 | 0.05324805 | 0.69082 | 0.4897 | 0.03637827 |
| Longitude | 1038081 | 0.1022 | -0.1232504 | -1.6799 | 0.09298 | -0.08846514 |

**Table 3:**
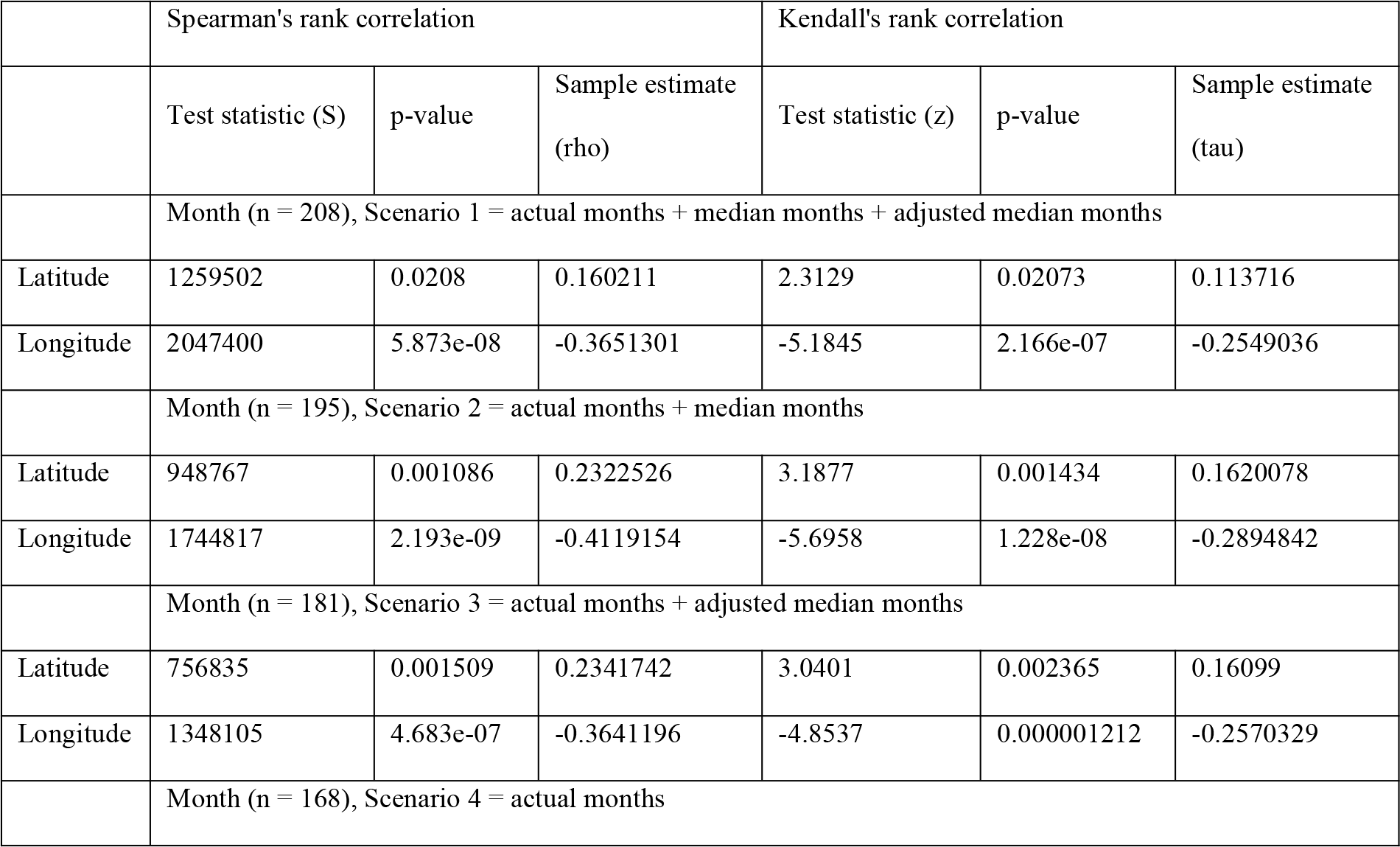
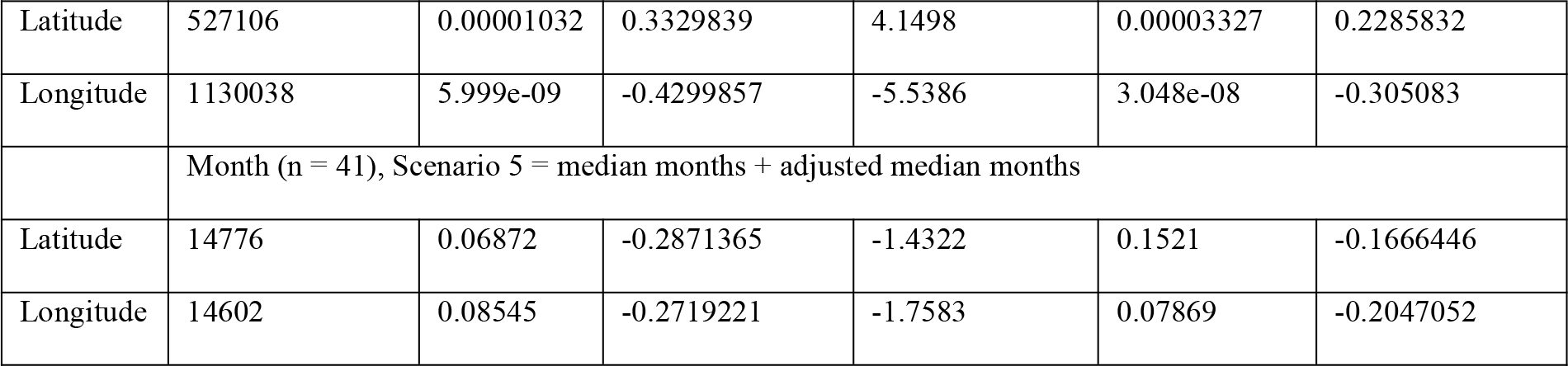
Correlations between months and coordinates.

**Table 4:**
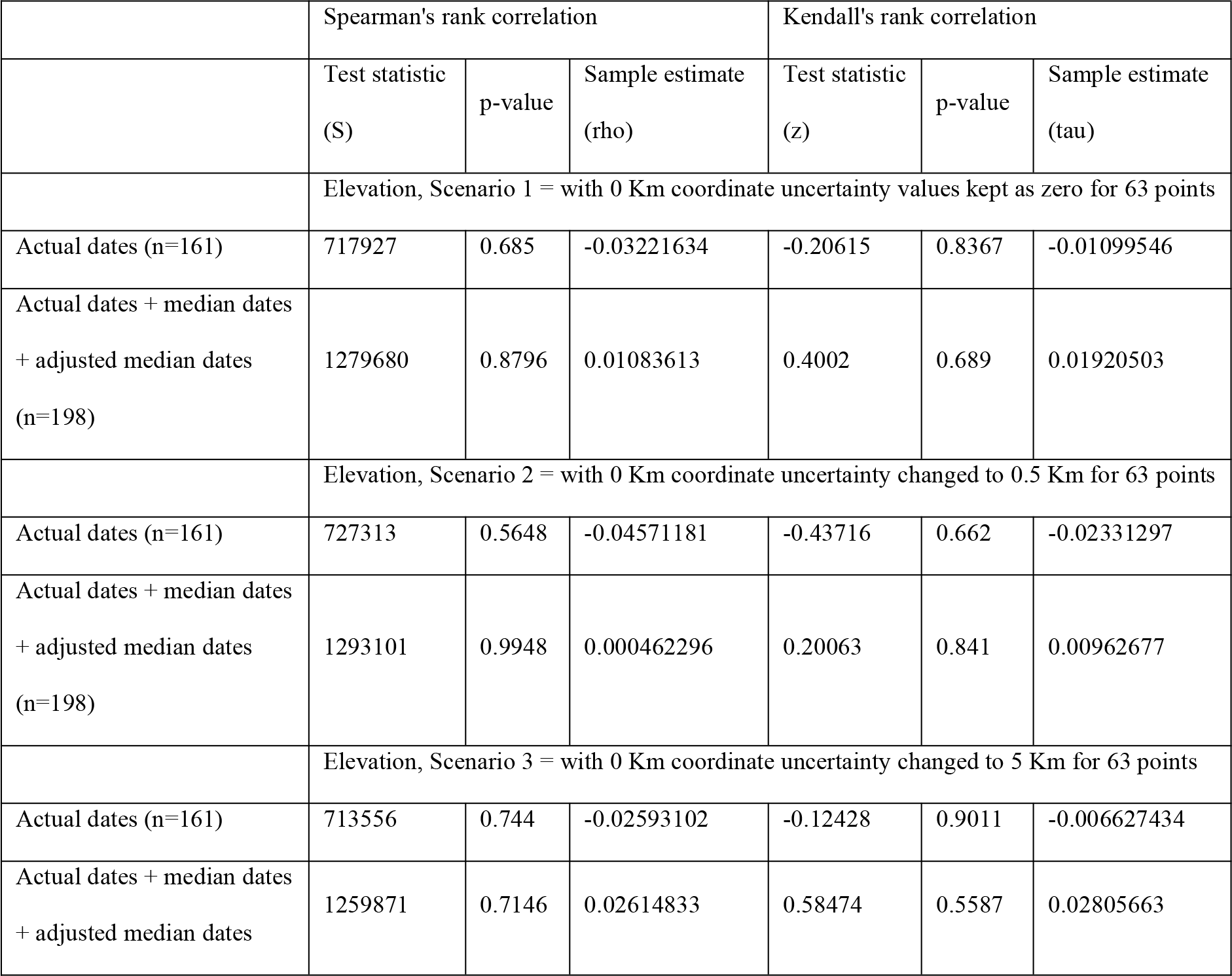
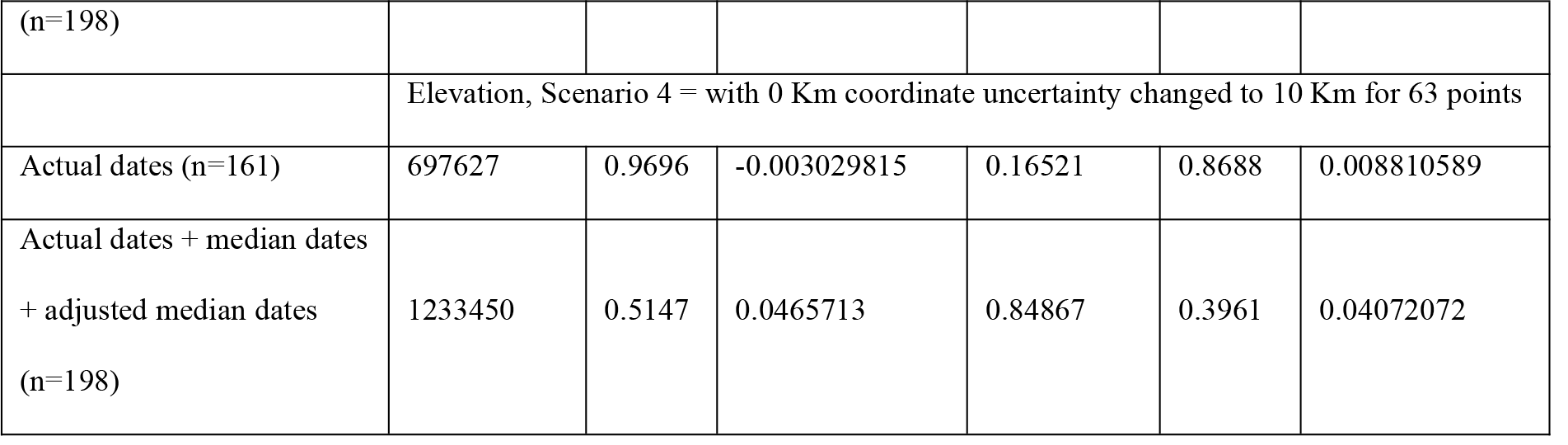
Correlations between elevations and dates.

**Table 5:** Correlations between elevations and coordinates.

|  | Spearman's rank correlation |  |  | Kendall's rank correlation |  |  |
| --- | --- | --- | --- | --- | --- | --- |
|  | Test statistic<br>(S) | p-value | Sample estimate<br>(rho) | Test statistic<br>(z) | p-value | Sample estimate<br>(tau) |
|  | Elevation (n = 216), Scenario 1 = with 0 Km coordinate uncertainty values kept as zero for 63 points |  |  |  |  |  |
| Latitude | 1216277 | 0.00003947 | 0.2758446 | 4.1632 | 0.00003138 | 0.1911641 |
| Longitude | 2031022 | 0.00199 | -0.2092438 | -2.7739 | 0.005539 | -0.1273734 |
|  | Elevation (n = 216), Scenario 2 = with 0 Km coordinate uncertainty changed to 0.5 Km for 63 points |  |  |  |  |  |
| Latitude | 1208391 | 0.00002871 | 0.2805397 | 4.377 | 0.00001203 | 0.2009557 |
| Longitude | 1967936 | 0.01149 | -0.1716836 | -2.1965 | 0.02806 | -0.1008476 |
|  | Elevation (n = 216), Scenario 3 = with 0 Km coordinate uncertainty changed to 5 Km for 63 points |  |  |  |  |  |
| Latitude | 1175178 | 0.000007048 | 0.3003145 | 4.4241 | 0.000009684 | 0.2031179 |
| Longitude | 1987323 | 0.006931 | -0.1832264 | -2.4621 | 0.01381 | -0.1130427 |
|  | Elevation (n = 216), Scenario 4 = with 0 Km coordinate uncertainty changed to 5 Km for 63 points |  |  |  |  |  |
| Latitude | 1155782 | 0.000002955 | 0.3118626 | 4.3864 | 0.00001152 | 0.2013881 |
| Longitude | 1974993 | 0.009592 | -0.1758851 | -2.3566 | 0.01844 | -0.1081993 |

**Table 6:** Correlations of elevations with and without buffer corrections.

|  | Test statistic (t) | Df | p-value | 95% CI | Sample estimate (PPMC) |
| --- | --- | --- | --- | --- | --- |
| Elevation, Scenario 1 = with 0 Km coordinate uncertainty values kept as zero for 63 points |  |  |  |  |  |
| Elevation without buffer correction (n=216) | 37.24 | 214 | < 2.2e-16 | 0.9103866 - 0.9466366 | 0.9307636 |
| Elevation without buffer correction (n=63) | Inf | 61 | < 2.2e-16 | 1 - 1 | 1 |
| Elevation, Scenario 2 = with 0 Km coordinate uncertainty changed to 0.5 Km for 63 points |  |  |  |  |  |
| Elevation without buffer correction (n=216) | 28.828 | 214 | < 2.2e-16 | 0.8607228 - 0.9161812 | 0.8917529 |
| Elevation without buffer correction (n=63) | 12.39 | 61 | < 2.2e-16 | 0.7568081 - 0.9041972 | 0.8459488 |
| Elevation, Scenario 3 = with 0 Km coordinate uncertainty changed to 5 Km for 63 points |  |  |  |  |  |
| Elevation without buffer correction (n=216) | 27.126 | 214 | < 2.2e-16 | 0.8460870 - 0.9070821 | 0.8801679 |
| Elevation without buffer correction (n=63) | 10.344 | 61 | 4.82e-15 | 0.68592 - 0.87317 | 0.7980576 |
| Elevation, Scenario 4 = with 0 Km coordinate uncertainty changed to 10 Km for 63 points |  |  |  |  |  |
| Elevation without buffer correction (n=216) | 24.521 | 214 | < 2.2e-16 | 0.8192044 - 0.8902195 | 0.8587826 |
| Elevation without buffer correction (n=63) | 7.9218 | 61 | 6.00e-11 | 0.5638130 - 0.8159125 | 0.7121022 |

Using one-way ANOVA, it was found that under two of the aforementioned conditions (n = 216 & n = 63), there was no significant difference in means among the four scenarios of elevation (p = 0.969 & p = 0.797 respectively). The results are visualised in Fig. 6 and enumerated in Table S3. In pairwise comparison of four scenarios of elevations, the value when n = 216, was 0.96 for all, and value when n = 63, was 0.92 for all. Bartlett’s test indicated that the variances were homogeneous in both cases (n = 216, Bartlett’s K-squared = 0.37976, df = 3, p-value = 0.9444 & n = 63, Bartlett’s K-squared = 1.0746, df = 3, p-value = 0.7832). There was also no discernible difference of means between latitudes and longitudes from the two groups (see Fig. S7).

**Figure 6:**
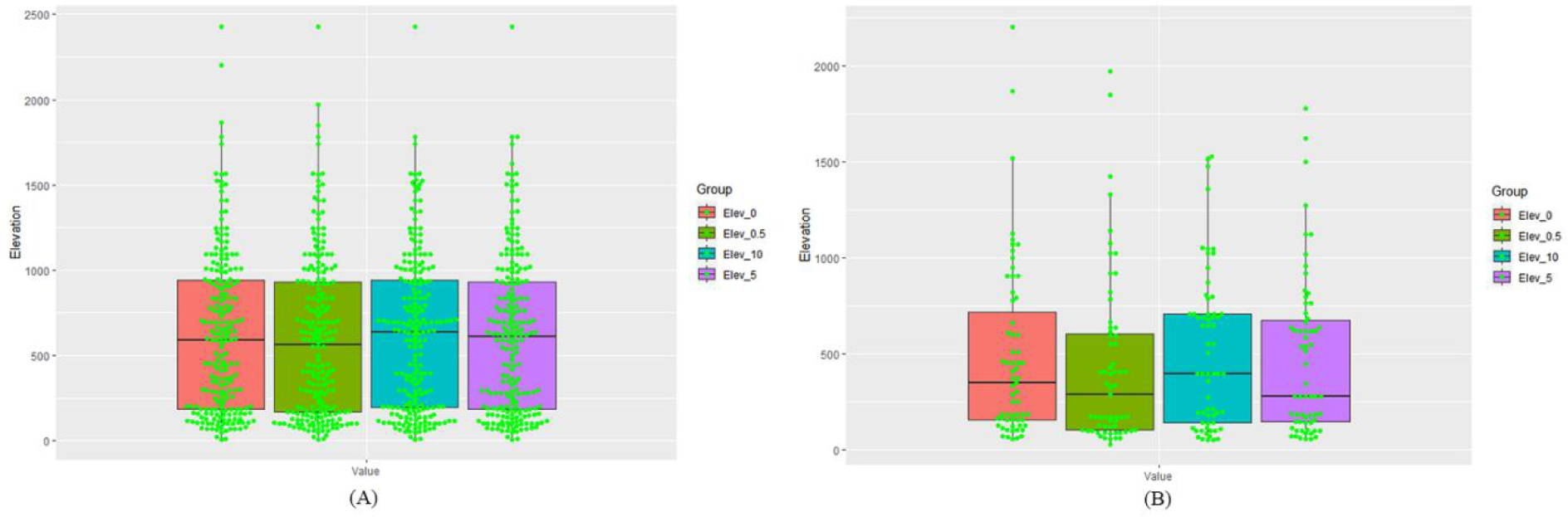
Comparison of spread of elevations under different scenarios. The four scenarios here are Elev 0, Elev 0.5, Elev 5, Elev 10 (with 0 km coordinate uncertainty kept as zero or changed to 0.5, 5, and 10 km respectively, for 63 points). (A) n = 216. (B) n = 63 (all points with varying buffer).

## Discussion

The sightings at Bilaspur district are the first records of the species/subspecies in this district, thereby filling a void in distribution along the western end of their range. They constitute the bulk of records from Himachal Pradesh (with only three pre-existing records from the state). If the westward expansion of the subspecies continues, sightings from the Hindukush mountain range of northern Pakistan and north-eastern Afghanistan can be expected, after which they will not able to survive due to the presence of dry deserts all around. It is unlikely that they will climb up the Himalayan range to higher elevations, as they prefer lower altitudes (this study provides partial evidence of this species’ resilience to climate change).

Nor are they expected to cross the Himalayan peaks, which acts as tall barriers for dispersal, a prediction affirmed by the absence of records from the northern side of the Himalayan range.

Studies using c-science data (primarily or along with other data) have found climate-induced expansions to (as well as retreat from) the northern latitudes in areas such as North America (Breed, Stichter & Crone, 2012) and northern Europe (Pöyry et al, 2009, Hällfors et al, 2021). The current study doesn’t provide any evidence for such a range shift but an in-depth study into the population biology could reveal more about this phenomenon. It could be that the numbers of this subspecies are growing in the northern portion of their range compared to stable populations in the southern portion.

In terms of seasonal variation, no significant linear patterns were observed, although Spearman’s correlation between longitude and months showed a moderate negative relationship. If this were to be a true correlation, then it would imply that as we move east, the activity of species is expected to peak earlier in the year. Histogram of months (Fig. 4C) shows peak activity in March and August-December, indicating annual bimodal distribution, with more activity in spring/post-winter, monsoon/pre-winter and winter/post-monsoon. This pattern may vary across their range. Due to the data limitations in some parts of their range, and the absence of long-term data, it is difficult to understand if the species has advanced their phenology as a response to climatic changes, as has been observed for a large number of butterflies in Finland (Hällfors et al, 2021), particularly relevant to less mobile or movement-restricted species (Amano et al, 2014, Socolar et al, 2017) like V. e.sinha.

When looking at seasonal patterns, one must look at the detection and non-detection data. Boruah et al (2018) reports non-detection during summer and monsoon months (March-October) and detection only during winter (November-February) in the Manchabandha Reserve Forest, Odisha. Similarly, non-detection was noted in the Barail Wildlife Sanctuary (Assam), for the months April-November and detection from December-March (Gogoi, Singha & Deb, 2016). In Tipong (eastern Assam), they were detected in September and not detected during the other months surveyed, representative of most seasons (Singh, 2017). In Pakke and Sessa in Arunachal Pradesh, they have been detected during May and September-November, and not detected during the other months surveyed (Sondhi & Kunte, 2016). Non-detection in summer and detection in winter, at the Baghmara Reserve Forest (Meghalaya), is reported by Kunte et al (2012). In the Santosh river catchment (Bhutan), non-detection was in March, May and July, whereas detection was in November and January. In Uttarakhand too, a similar pattern is seen — detection in March, and September-December, and non-detection during the rest of the year (Singh & Sondhi, 2016; Sondhi, 2021, pers. obs.). Smetacek (2012) reports activity in May and July-October (and hence bins them under the multivoltine category which is used to refer to species that usually have multiple overlapping generations). This is also a little different from the current study (August-November). From all this, it can be inferred that they are more active during the winter months, at least in and near the North- East Indian cluster and the Western Himalayan cluster.

Butterfly species, especially mountain butterflies have been reported to climb up to the summits over the years as a result of climate change (Molina-Martínez et al, 2016; Rödder et al, 2021). No patterns associated with elevation were observed for V. e. sinha. This means that the species has not climbed up over the years as a result of climate change. Molina-Martínez et al (2016) reported that the species occurring below the elevation of 1000 m, as well as all the species combined, had significantly less range shifts towards higher elevations than what was expected based only on global warming. Our results are consistent with this finding because V. e. sinha occurs mostly below 1000 m. This could either be as a result of their resilience as a species or their annual life cycle where they are more active during the colder part of the year. The populations seems to be randomly spread across the elevational gradient across their range of distribution. This could alternatively be a complex pattern which is yet to be understood.

Mean distance between the sets of points (∼7.85 km), is higher than the known daily dispersal of the species — around 1-5 km (Smetacek, 2012). This can negatively affect studies that require a high resolution of information such as studies on microhabitats of the species. This difference is unlikely to affect research such as Species Distribution Modelling (Wisz et al, 2008; Mizsei et al, 2016) because of high performance even with low spatial accuracy and lower number of points for models such as MaxEnt. As is seen in Fig. 6b, when using just the coordinates with unknown uncertainty as opposed to using them when mixed with coordinates with fixed buffers (Fig. 6a), there is a notable variation in means, which can become significant when the dataset is largely or wholly the former type of coordinates. So, metadata associated with datapoints in c-science datasets can affect the results. For this reason, biological plausibility and relevance should be considered while deciding which datum to keep and which to exclude. Some butterflies are known to be high-flyers and others low-flyers (DeVries, Penz & Hill, 2010). The former may use canopies to move around which are parallel to the slopes in hilly areas which translates into wide variation in elevation, whereas the latter may use roads which are generally along the contours or slopes with a low gradient of elevation. So, averaging elevation using a buffer may have different effects as covariates on different species. For proper use and application of data (usually from c-science sources), one must have some idea about the behaviour and movement ecology of the species under study.

The problems associated with the collection and use of data, and potential remedies, are elucidated in Table 7.

**Table 7:**
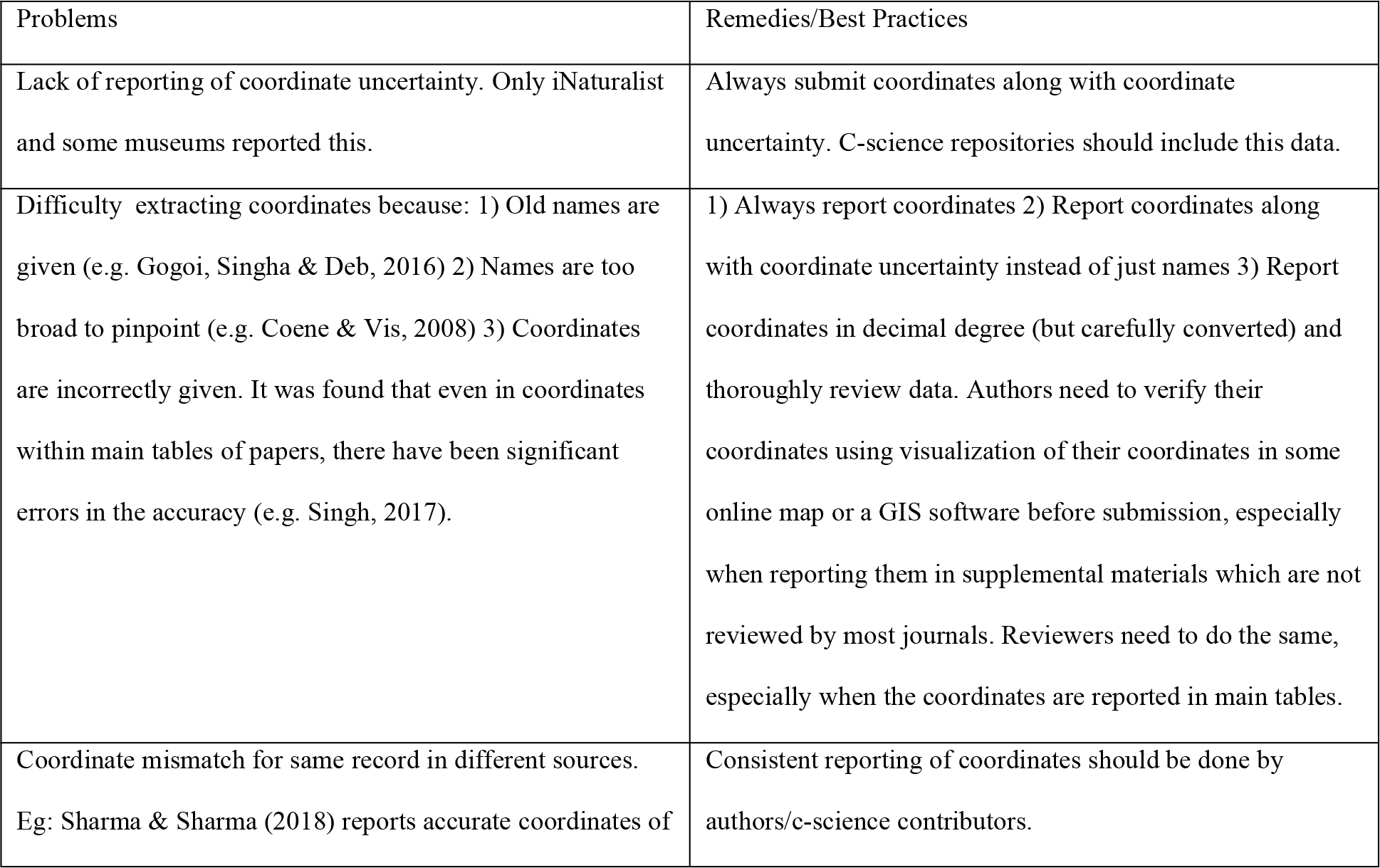
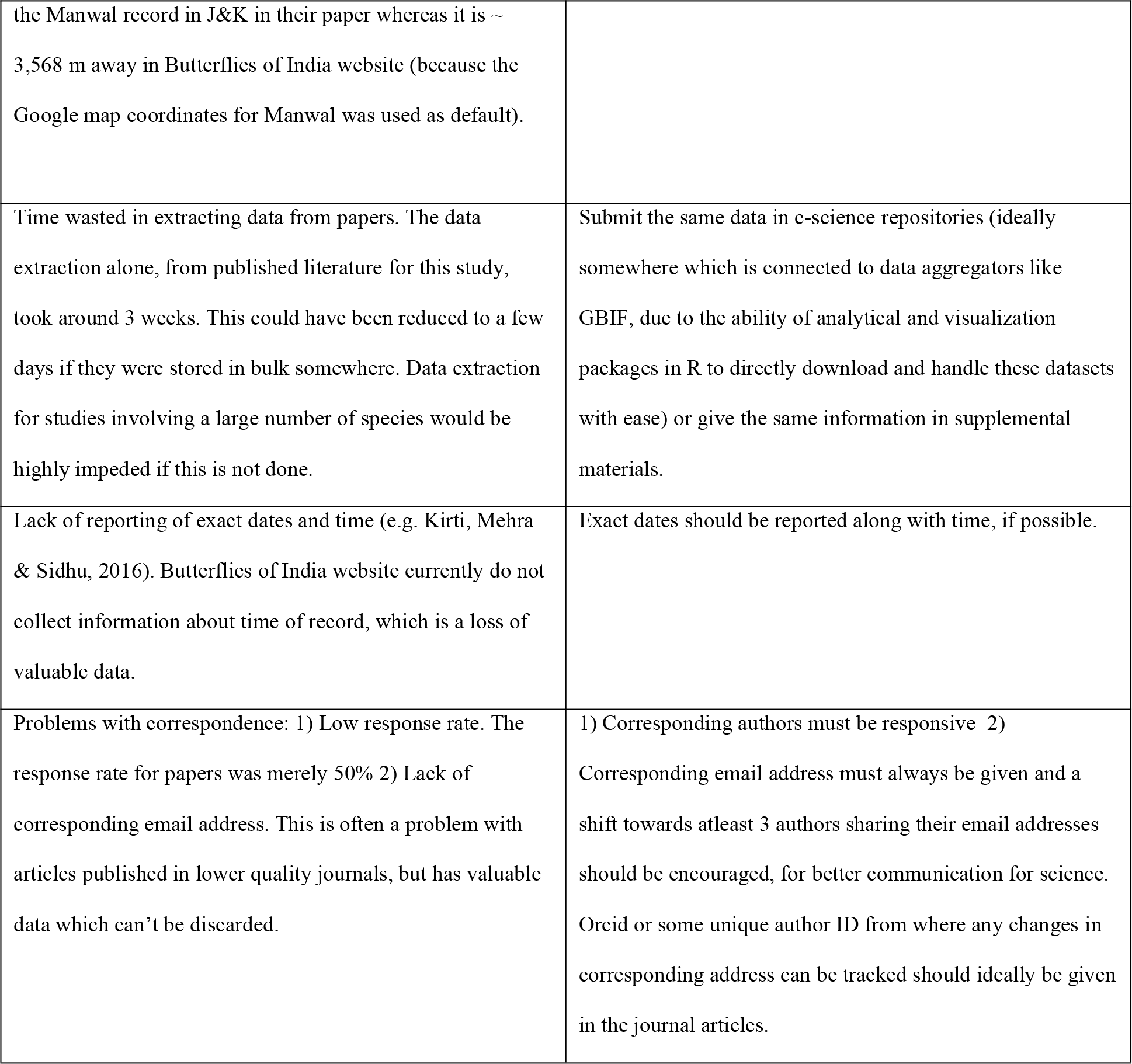
Problems with, and remedies/best practices for data collection and use.

## Conclusion

In the current study, it has been found that V. e. sinha has likely undergone a range expansion. None of the other potential spatiotemporal phenomena that were investigated, exhibited any discernible patterns. The current study has helped in adding to the knowledge on this subspecies, such as their distribution patterns and seasonal activity. It has brought more private data into the public domain (Supplemental Table S4 contain the raw and derived data used for the research). This data can be used for some more computationally expensive analyses such as maximum entropy modelling which can address the some of the questions asked in the current study with more accuracy. It has identified knowledge gaps which other researchers could look into. It has been found that we need more c-science data, especially from before 2000. This can be achieved by writing to older researchers and helping them to digitise the data which are gathering dust in their field notes, thus adding professionally-collected data to c-science repositories. As has been noticed in this study, we need more precise data. This would require teaching the c-science contributors about the best practices in data entry. To that extent, we would need better curation and peer-review of data, whether in journals or in c-science repositories. The results from the current study come with the caveat of the obvious sampling bias, with a high amount of sampling in much of North-East India and Uttarakhand compared to other regions. It was found that the way c-science data are extracted and handled can potentially affect the results of a study. As is evident from the varying taxonomy of the genus, species, and subspecies (briefly described in the introduction), this genus needs a taxonomic revision. Studies like this help in such an endeavour by providing ecological details that may be useful in taxonomic revisions based on multiple lines of evidence, including genetics (as has been carried out for the genus Cethosia Fabricius belonging to the same subfamily as Vagrans - Heliconiinae (Müller & Beheregaray, 2010)).

## Acknowledgements

We thank the Himachal Pradesh Forest Department for the permission granted to carry out our research in Bilaspur district. We acknowledge Dr. Krushnamegh Kunte for sharing the dataset of contributions to the Butterflies of India website. We thank Tshetsholo Naro, Bitupan Boruah and Dr. Monwar Hossain for providing details of records from their respective papers. Our gratitude goes towards Dr. Peter V. Küppers, Dr. Mutum Ingobi Singh (prompted by Dr. Ramaiyer Varatharajan), and Dr. Roger Clive Kendrick, for not only giving details of records from their respective publication, but other unpublished records of the species. We would also like to thank all the remaining contributors of the records through c- science, published literature and other sources. We would like to thank Faezal Yunus, Dr. Shankha Banerjee, Abhishek Hariharan and Benny Malone for proofreading and bettering this manuscript.

## Supplemental Information

**Suppmental Figure S1.**
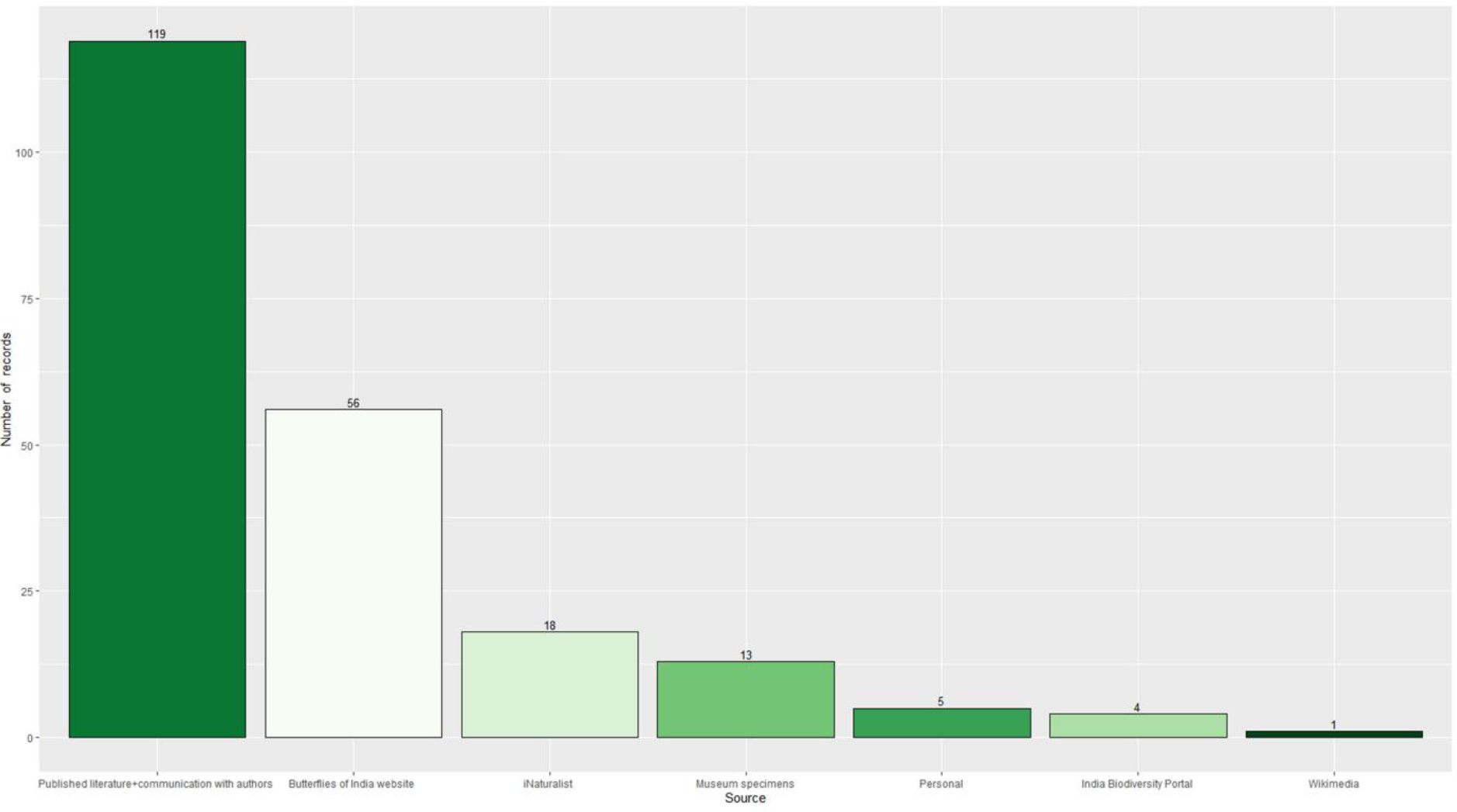
Contribution of records by sources.

**Suppmental Figure S2:**
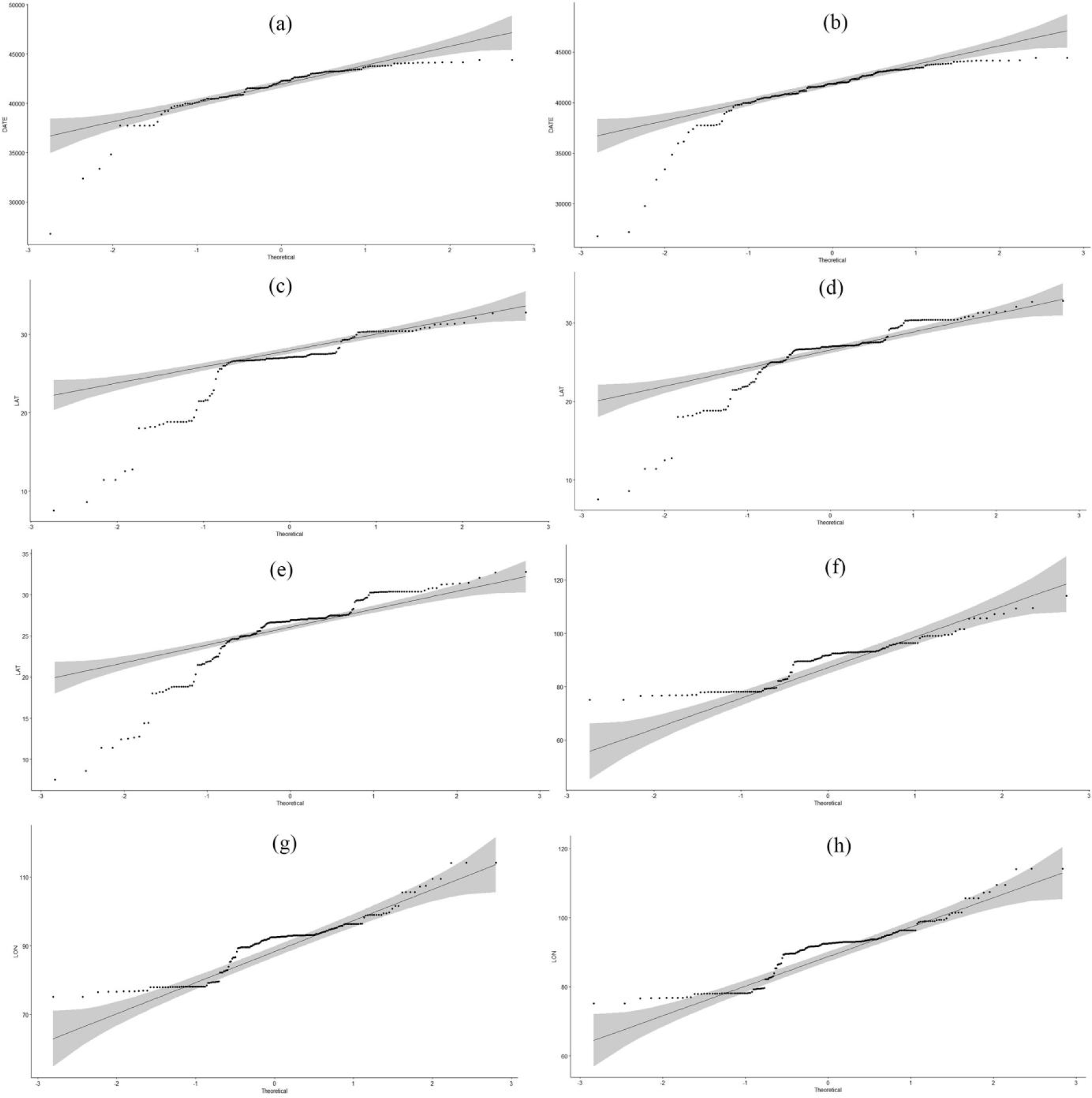
q-q plots for dates, latitudes, and longitudes. (a) Dates (n = 161, Scenario I = only actual dates), (b) Dates (n = 198, Scenario 2 = actual dates + median dates + adjusted median dates), (c) Latitudes (n = 161), Latitudes (n = 198), (e) Latitudes (n = 2 16), (f) Longitudes (n = 161), (g) Longitudes **(n** = 198), and (h) Longitudes (n = 2 16). Please note that only two or three examples are given for each here, but q-q plots were actually generated prior to each correlations. This isju stified for the following reasons - i) Once it ’s established that a set with highest number of observations is non-normal (e.g. longitudes with 2 16 observations), it can be inferred that normal ity tends to be non-normal for a smaller subset of it (e.g. longitudes with 33 observations), as there is more variability ii) Non-parametric tests were carried out even if one in a pair of variables is found to be non-normal.

**Suppmental Figure S3:**
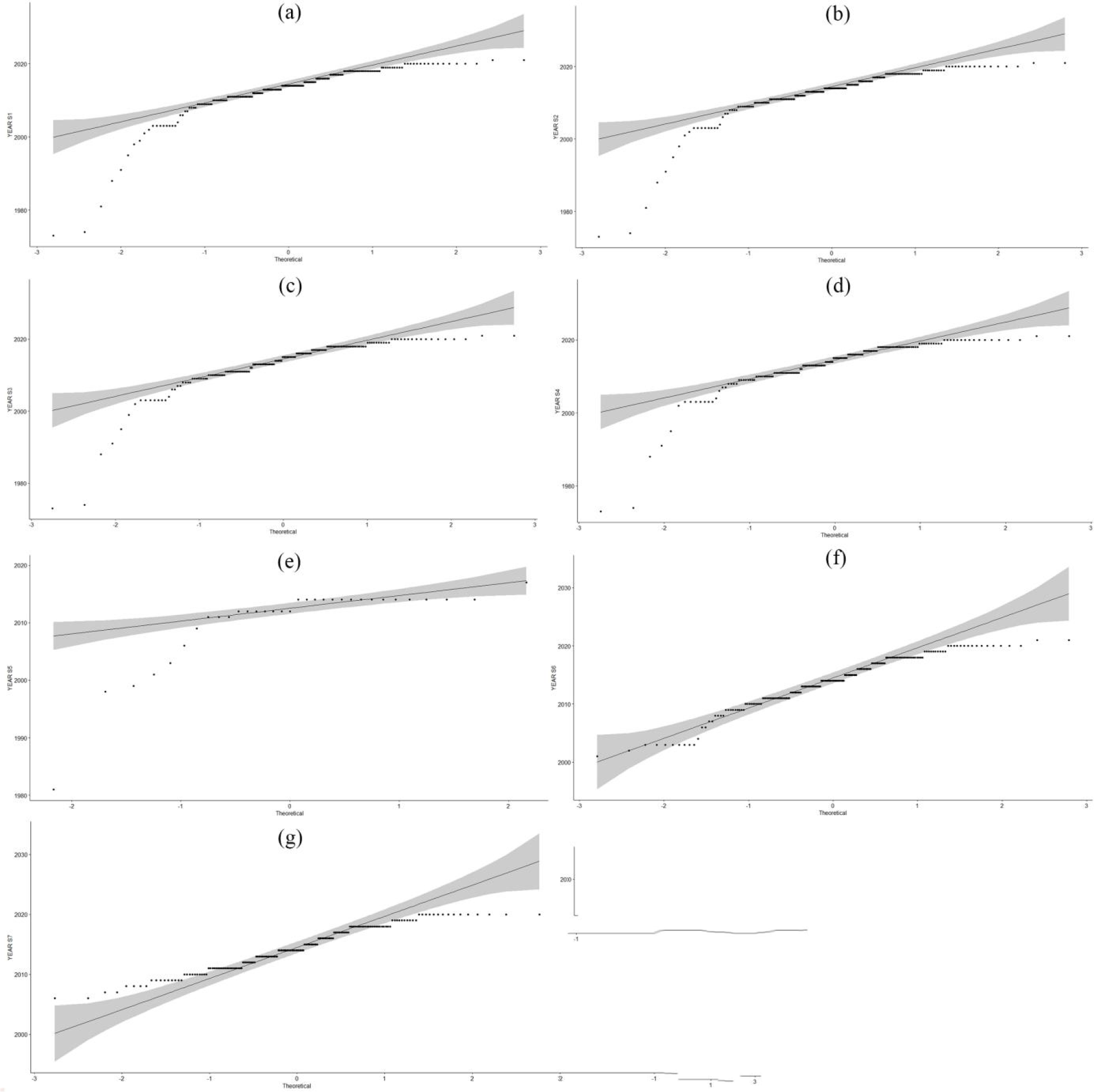
q-q plots for years. (a) n = 198, Scenario I = acn1al years + median years + adjusted med ian years, (b) n = 195, Scenario 2 = acnial years + median years, (c) n = 168, Scenario 3 = acn1al years + adjusted median years, (d) n = 165, Scenario 4 = acn1al years, (e) n = 33, Scenario 5 = median years + adju sted median years, (t) ) n = 190, Scenario 6 = Scenario I - pre-2000 records, and (g) n = 177, Scenario 7 = Scenario I - pre-2005 records.

**Suppmental Figure S4:**
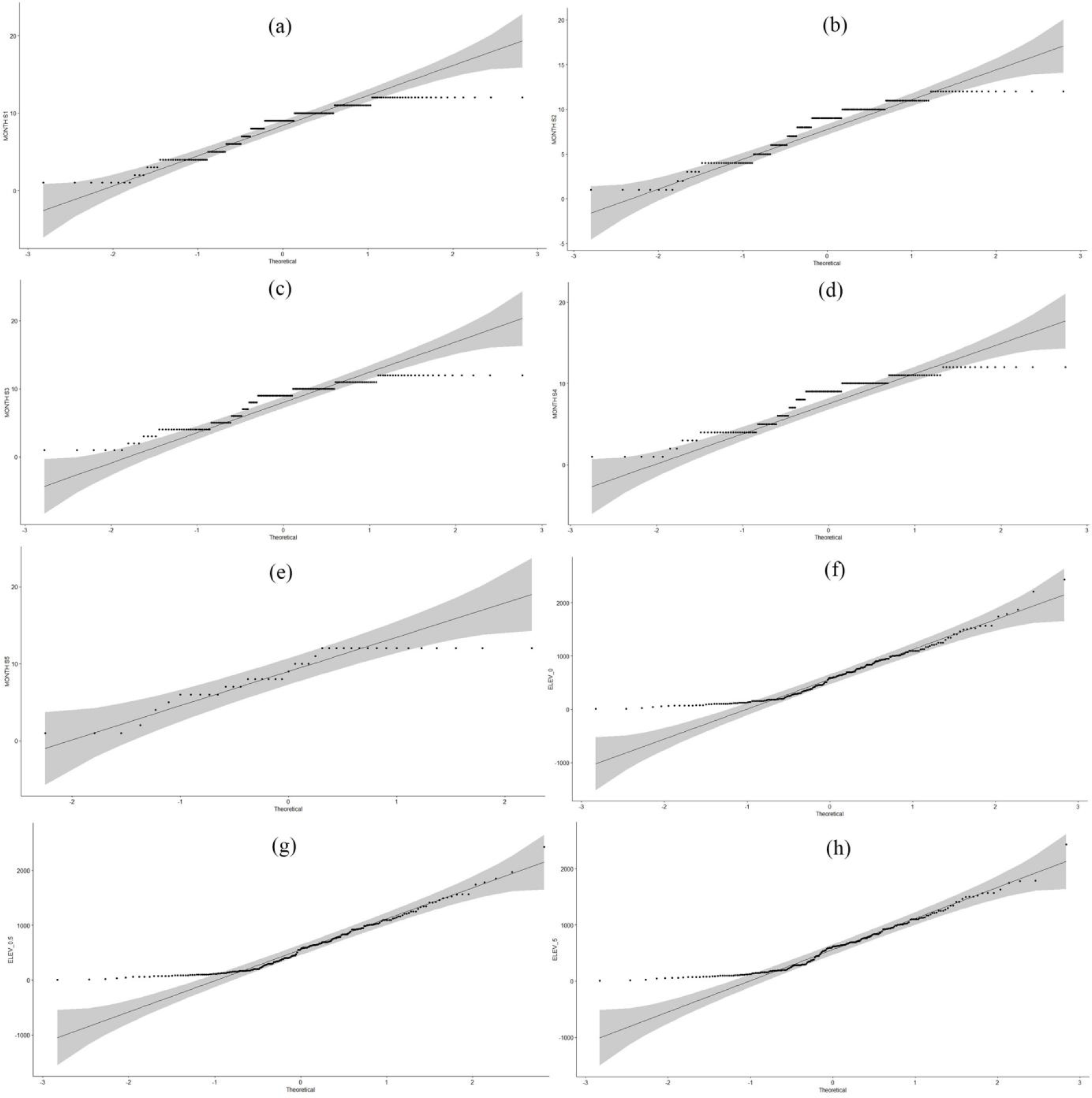
q-q plots for months and elevations. (a) Months (n = 208, Scenario I = actual months + median months + adjusted median months), (b) Months **(n** = 195, Scenario 2 = actual months + median months), (c) Months (n = 181, Scenario 3 = actual months + adju sted median months), (d) Months (n = 168, Scenario 4 = actual months), (d) Months (n = **41,** Scenario 5 = median months + adjusted median months), (f) Elevations (n = 216, Scenario I = with 0 Km coordinate uncertainity values kept as zero for 63 points),(g) Elevations (n = 2 16, Scenario 2 = with 0 Km coordinate uncertainity changed to 0.5 Km for 63 points), and (h) Elevations (n = 2 16, Scenario 3 = with 0 Km coordinate uncertainitv changed to 5 Km for 63 points).

**Suppmental Figure S5:**
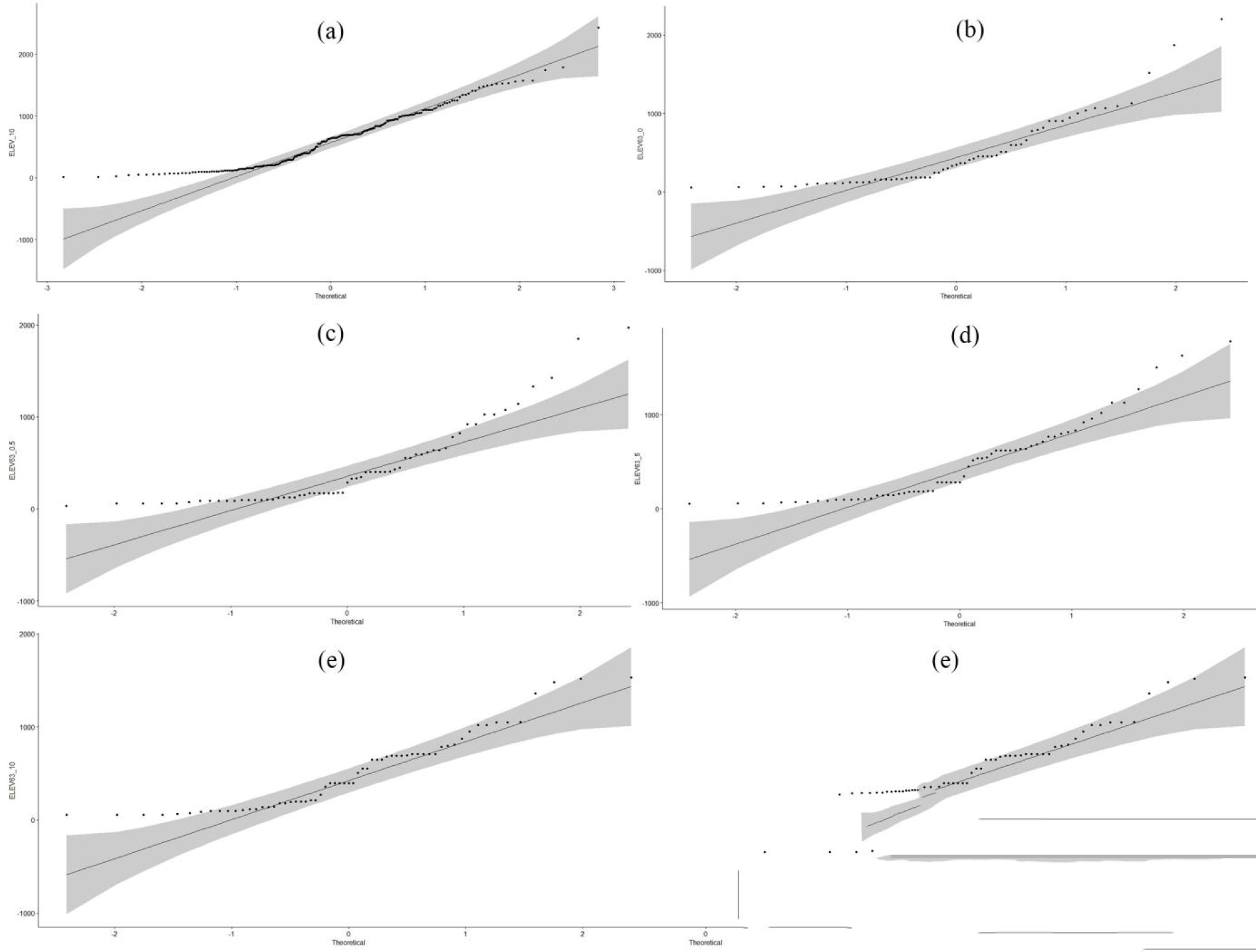
q-q plots for elevations. n = 2 16, Scenario 4 = with 0 Km coordinate uncerta inity changed to 10 Km for 63 points, (b) n = 63, Scenario I = with 0 Km coordinate uncertainity values kept as zero for all points, (c) n = 63, Scenario 2 = with 0 Km coordinate uncertainity changed to 0.5 Km for all points, (d) n = 63, Scenario 3 = with 0 Km coordinate uncertainity changed to 5 Km for all points, and (e) n = 63, Scenario 4 = with 0 Km coordinate uncertainity changed to 10 Km for all points.

**Supplemental Figure S6.**
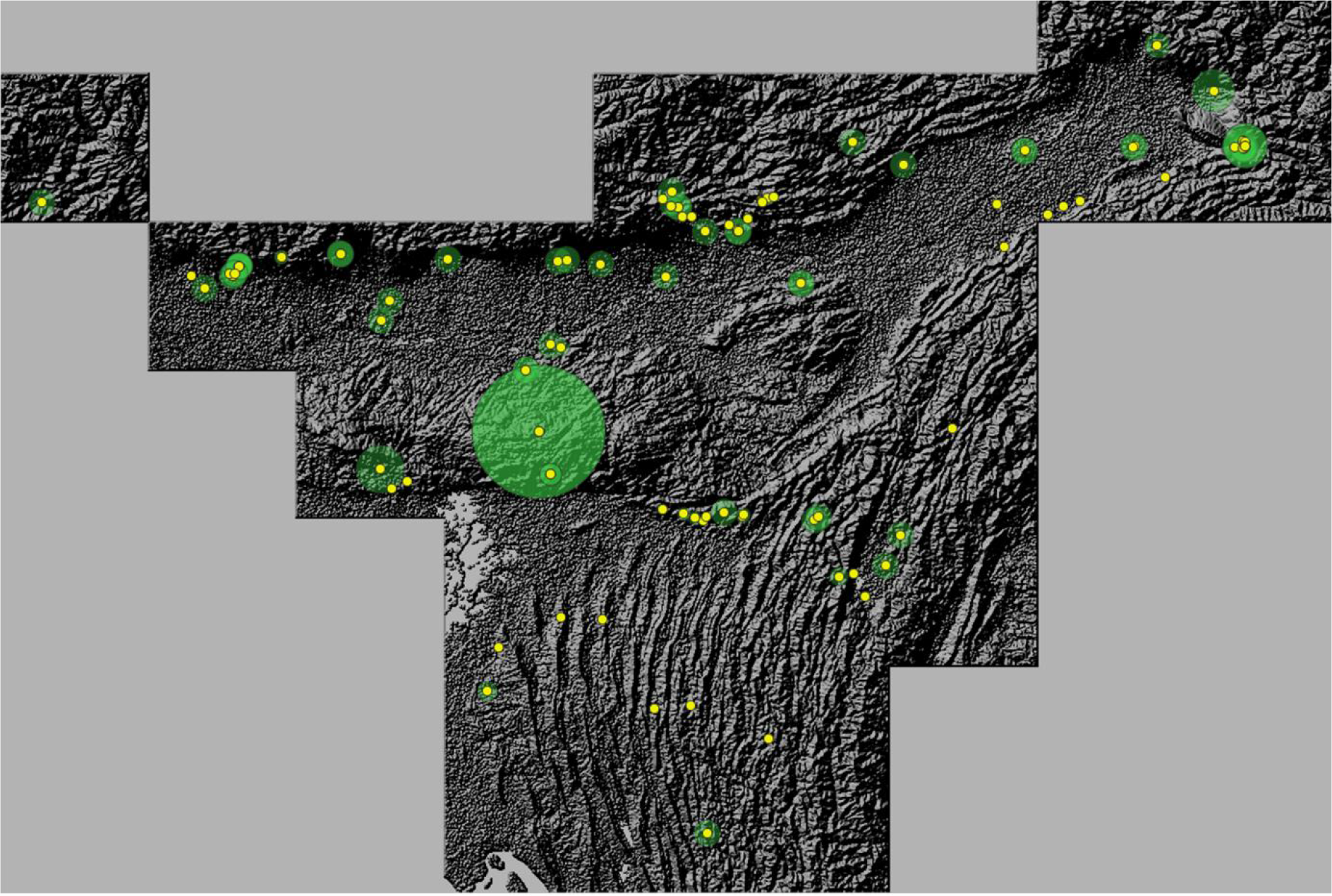
Buffers around coordinate points in North-East India cluster over hillshade map.

**Supplemental Figure S7.**
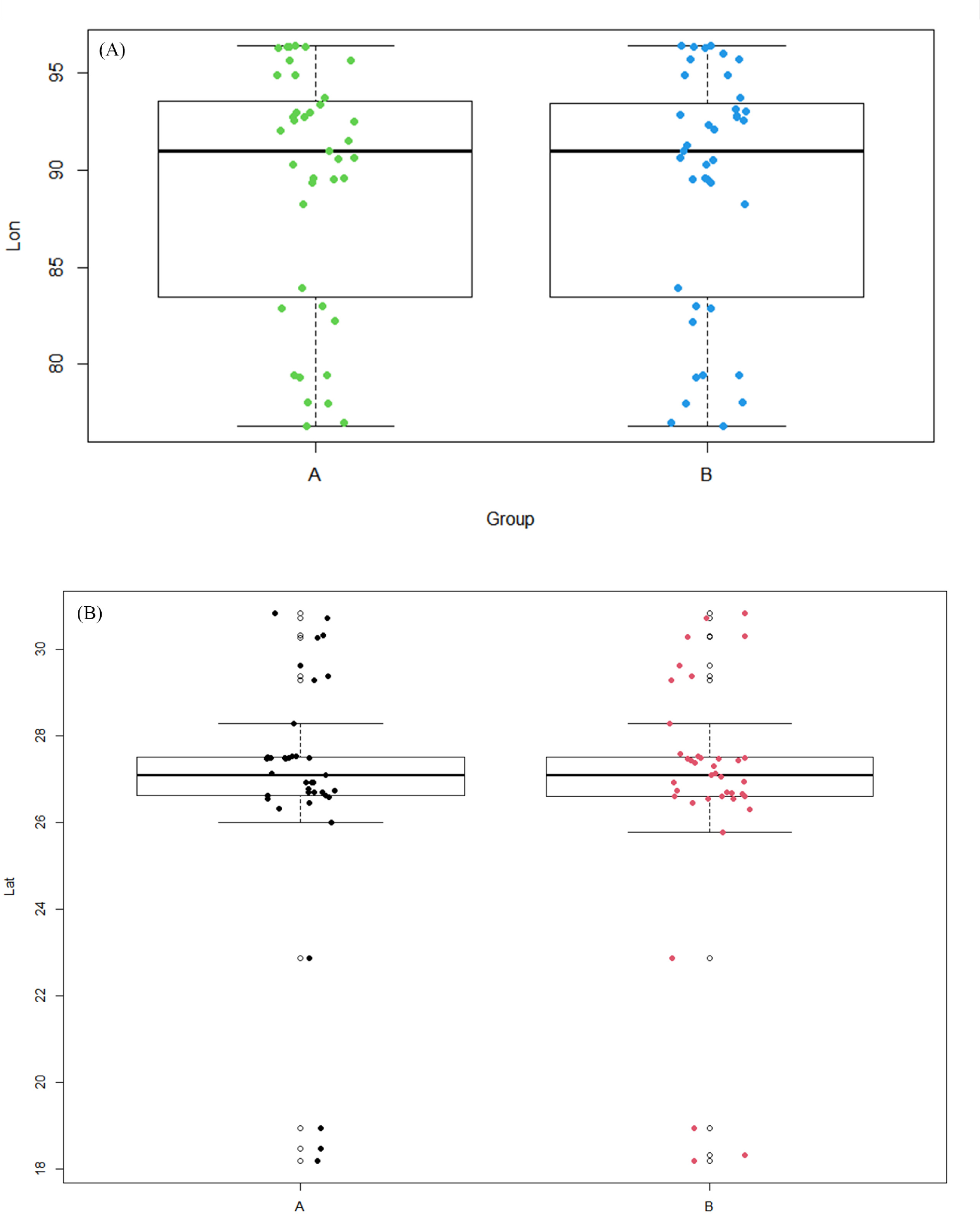
Comparison of spread of coordinates amongst two groups - derived from names and acquired through request. (A) Spread of longitudes. (B) Spread of latitudes.

**Supplemental Table S1.**
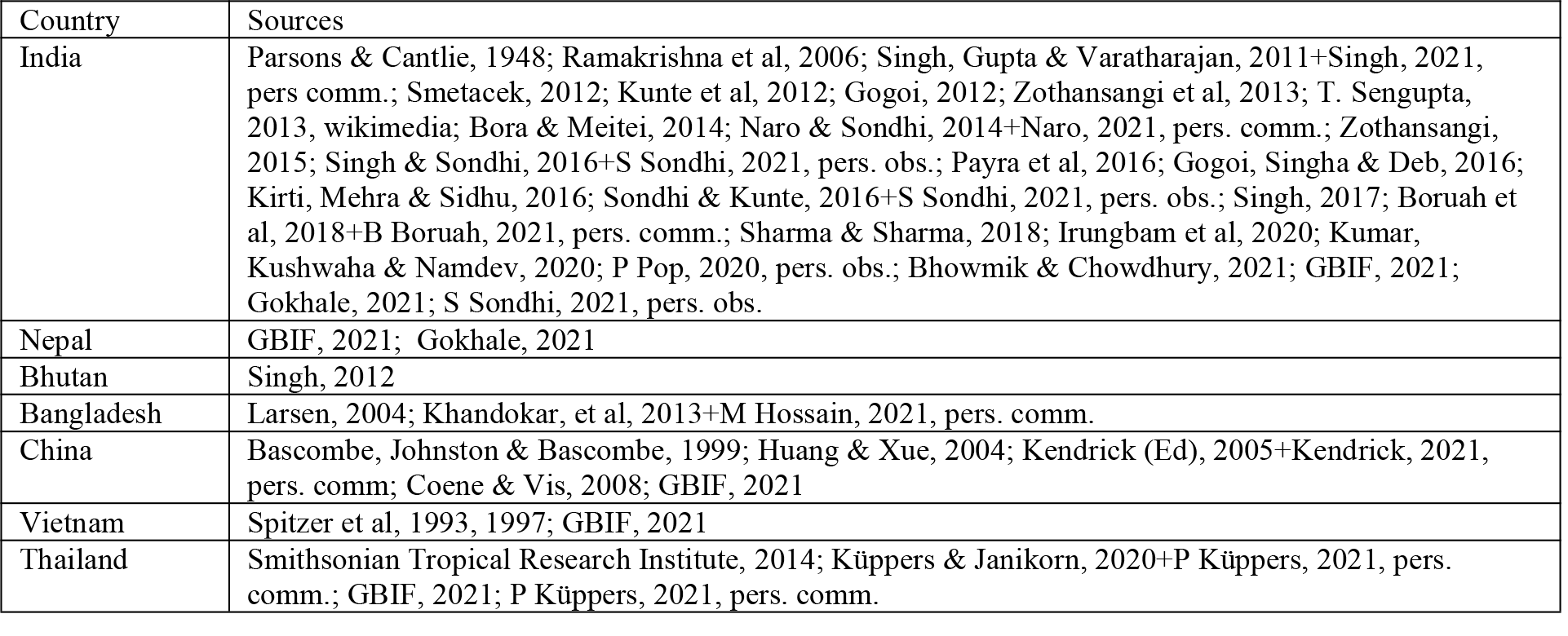
Sources of records for each country.

Sources of records for each country.
| Country | Sources |
| --- | --- |
| India | Parsons & Cantlie, 1948; Ramakrishna et al, 2006; Singh, Gupta & Varatharajan, 2011+Singh, 2021, pers comm.; Smetacek, 2012; Kunte et al, 2012; Gogoi, 2012; Zothansangi et al, 2013; T. Sengupta, 2013, wikimedia; Bora & Meitei, 2014; Naro & Sondhi, 2014+Naro, 2021, pers. comm.; Zothansangi, 2015; Singh & Sondhi, 2016+S Sondhi, 2021, pers. obs.; Payra et al, 2016; Gogoi, Singha & Deb, 2016; Kirti, Mehra & Sidhu, 2016; Sondhi & Kunte, 2016+S Sondhi, 2021, pers. obs.; Singh, 2017; Boruah et al, 2018+B Boruah, 2021, pers. comm.; Sharma & Sharma, 2018; Irungbam et al, 2020; Kumar, Kushwaha & Namdev, 2020; P Pop, 2020, pers. obs.; Bhowmik & Chowdhury, 2021; GBIF, 2021; Gokhale, 2021; S Sondhi, 2021, pers. obs. |
| Nepal | GBIF, 2021; Gokhale, 2021 |
| Bhutan | Singh, 2012 |
| Bangladesh | Larsen, 2004; Khandokar, et al, 2013+M Hossain, 2021, pers. comm. |
| China | Bascombe, Johnston & Bascombe, 1999; Huang & Xue, 2004; Kendrick (Ed), 2005+Kendrick, 2021, pers. comm; Coene & Vis, 2008; GBIF, 2021 |
| Vietnam | Spitzer et al, 1993, 1997; GBIF, 2021 |
| Thailand | Smithsonian Tropical Research Institute, 2014; Küppers & Janikorn, 2020+P Küppers, 2021, pers. comm.; GBIF, 2021; P Küppers, 2021, pers. comm. |

**Supplemental Table S2.** Shapiro-Wilk values of variables

| Supplemental Table S2 - Shapiro-Wilk values of variables |  |  |
| --- | --- | --- |
| Variable | W | p-value |
| Date (n = 161, Scenario 1 = only actual dates) | 0.79526 | 9.701E-14 |
| Date (n = 198, Scenario 2 = actual dates + median dates + adjusted median dates) | 0.76713 | < 2.2E-16 |
| Latitude (n = 161, corresponding to Dates - Scenario 1) | 0.80722 | 2.679E-13 |
| Latitude (n = 198, corresponding to Dates - Scenario 2) | 0.84975 | 4.969E-13 |
| Latitude (n = 216) | 0.86059 | 3.891E-13 |
| Longitude (n = 161, corresponding to Dates - Scenario 1) | 0.92683 | 0.00000027 |
| Longitude (n = 198, corresponding to Dates - Scenario 2) | 0.92254 | 1.025E-08 |
| Longitude (n = 216) | 0.92263 | 3.222E-09 |
| Year (n = 198, Scenario 1 = actual years + median years + adjusted median years) | 0.76388 | < 2.2E-16 |
| Year (n = 195, Scenario 2 = actual years + median years) | 0.75486 | < 2.2E-16 |
| Year (n = 168, Scenario 3 = actual years + adjusted median years) | 0.76362 | 3.616E-15 |
| Year (n = 165, Scenario 4 = actual years) | 0.75303 | 2.297E-15 |
| Year (n = 33, Scenario 5 = median years + adjusted median years) | 0.63378 | 7.453E-08 |
| Year (n = 190, Scenario 6 = Scenario 1 - pre-2000 records) | 0.94735 | 0.000001851 |
| Year (n = 177, Scenario 7 = Scenario 1 - pre-2005 records) | 0.95856 | 0.00004373 |
| Month (n = 208, Scenario 1 = actual months + median months + adjusted median months) | 0.90442 | 2.732E-10 |
| Month (n = 195, Scenario 2 = actual months + median months) | 0.90965 | 1.547E-09 |
| Month (n = 181, Scenario 3 = actual months + adjusted median months) | 0.89125 | 3.196E-10 |
| Month (n = 168, Scenario 4 = actual months) | 0.89414 | 1.354E-09 |
| Month (n = 41, Scenario 5 = median months + adjusted median months) | 0.84783 | 0.0000663 |
| Elevation (n = 216, Scenario 1 = with 0 Km coordinate uncertainty values kept as zero for 63 points) | 0.92148 | 2.625E-09 |
| Elevation (n = 216, Scenario 2 = with 0 Km coordinate uncertainty changed to 0.5 Km for 63 points) | 0.91845 | 1.549E-09 |
| Elevation (n = 216, Scenario 3 = with 0 Km coordinate uncertainty changed to 5 Km for 63 points) | 0.92817 | 8.846E-09 |
| Elevation (n = 216, Scenario 4 = with 0 Km coordinate uncertainty changed to 10 Km for 63 points) | 0.9345 | 2.975E-08 |
| Elevation (n = 63, Scenario 1 = with 0 Km coordinate uncertainty values kept as zero for all points) | 0.81991 | 2.641E-07 |
| Elevation (n = 63, Scenario 2 = with 0 Km coordinate uncertainty changed to 0.5 Km for all points) | 0.79862 | 7.311E-08 |
| Elevation (n = 63, Scenario 3 = with 0 Km coordinate uncertainty changed to 5 Km for all points) | 0.86418 | 0.000005268 |
| Elevation (n = 63, Scenario 4 = with 0 Km coordinate uncertainty changed to 10 Km for all points) | 0.8964 | 0.00006658 |

**Supplemental Table S3.**
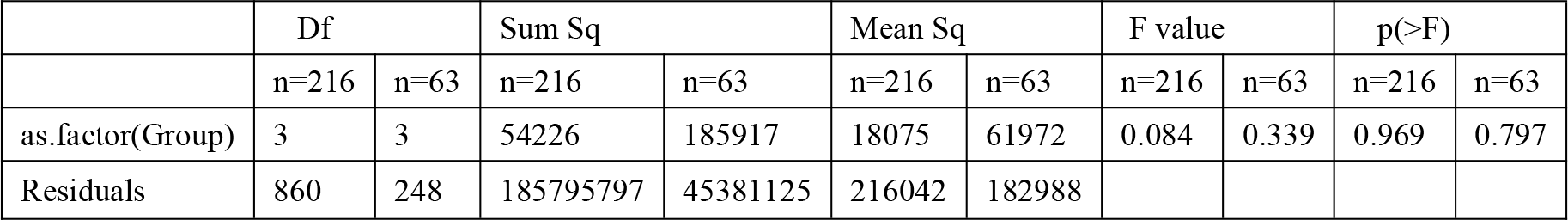
One-way ANOVA for testing difference in means between elevations under different scenarios.

|  | Df |  | Sum Sq |  | Mean Sq |  | F value |  | p(>F) |  |
| --- | --- | --- | --- | --- | --- | --- | --- | --- | --- | --- |
|  | n=216 | n=63 | n=216 | n=63 | n=216 | n=63 | n=216 | n=63 | n=216 | n=63 |
| as.factor(Group) | 3 | 3 | 54226 | 185917 | 18075 | 61972 | 0.084 | 0.339 | 0.969 | 0.797 |
| Residuals | 860 | 248 | 185795797 | 45381125 | 216042 | 182988 |  |  |  |  |

Supplemental Table S4: Raw and derived data

Mendeley Data, V1, doi: 10.17632/mpn65fcnwb.1 (All other supplementary materials included in the text can also be found here)

## Notes

https://doi.org/10.17632/mpn65fcnwb.1

### Competing Interest Statement

The authors have declared no competing interest.

